# A Vulnerable Subtype of Dopaminergic Neurons Drives Early Motor Deficits in Parkinson’s Disease

**DOI:** 10.1101/2024.12.20.629776

**Authors:** Akira Fushiki, David Ng, Zachary R. Lewis, Archana Yadav, Tatiana Saraiva, Luke A. Hammond, Christoph Wirblich, Bosiljka Tasic, Vilas Menon, Joaquim Alves da Silva, Rui M. Costa

**Affiliations:** Allen Institute, Seattle, WA 98109, USA; Zuckerman Mind Brain Behavior Institute, Columbia University, New York, NY 10027, USA; Center for Translational and Computational Neuroimmunology, Department of Neurology Columbia University Irving Medical Center, New York, NY 10032, USA; Champalimaud Research, Champalimaud Foundation, Lisbon 1400-038, Portugal; Department of Neurology, University Hospital of Würzburg, Würzburg 97080, Germany; Department of Neurology, The Ohio State University, Columbus, OH 43210, USA; Department of Microbiology and Immunology, Sidney Kimmel Medical College, Thomas Jefferson University, Philadelphia, PA 19107, USA; NOVA Medical School, Universidade Nova de Lisboa, Lisbon 1169-056, Portugal; Aligning Science Across Parkinson’s (ASAP) Collaborative Research Network, Chevy Chase, MD, 20815

**Keywords:** Parkinson’s disease, dopamine, cell types, neurodegeneration, vulnerability, tremor, motor symptoms

## Abstract

In Parkinson’s disease (PD), dopaminergic neurons (DANs) in the midbrain gradually degenerate, with ventral substantia nigra pars compacta (SNc) DANs exhibiting greater vulnerability. However, it remains unclear whether specific molecular subtypes of ventral SNc DANs are more susceptible to degeneration in PD, and if they contribute to the early motor symptoms associated with the disease. We identified a subtype of *Sox6+* DANs, *Anxa1+*, which are selectively lost earlier than other DANs, and with a time course that aligns with the development of motor symptoms in MitoPark mice. We generated a knock-in Cre mouse line for *Anxa1+* DANs and showed differential anatomical inputs and outputs of this population. Further, we found that the inhibition of transmitter release in *Anxa1+* neurons led to bradykinesia and tremor. Therefore, *Anxa1+* is not only a biomarker of a selectively vulnerable subtype of DANs, but is also sufficient to drive early motor symptoms in Parkinson’s disease.

## Introduction

Parkinson’s disease (PD) is a progressive neurological disorder that primarily affects movement. It develops gradually, sometimes starting with a barely noticeable tremor in one hand^1^. While tremors, rigidity and akinesia are well-known signs of PD, the most distinctive symptom is bradykinesia, characterized by the slowing of movements. The main pathological hallmark of PD is the loss of dopaminergic neurons (DANs) in the substantia nigra^2,3^ and the progressive reduction of DANs worsens the motor symptoms over time.

DANs in the midbrain are heterogeneous and they differ in their cellular properties and functions^4,5^. This heterogeneity is evident across several dimensions, including their molecular profiles and connectivity^4,6^. Transcriptomic studies have revealed that populations of midbrain DANs can be differentiated by distinct gene expression profiles^4^, a finding that extends beyond the conventional anatomical division of midbrain DANs into the substantia nigra pars compacta (SNc, A9) and the ventral tegmental area (VTA, A10). Recent findings have discovered that molecularly unique subtypes of DANs innervate different subregions of the striatum, suggesting a basis for the variety of behavioral responses linked to these distinct neural pathways^5,6^. In PD, the degeneration of DANs follows a stereotypical pattern where some populations of neurons are more vulnerable than others^7^. Within the SNc, there is a distinction between the ventral-tier and dorsal-tier DANs, where the ventral-tier neurons tend to degenerate first^7^. Other studies have focused on changes in gene expression associated with PD compared to healthy controls^8^. Transcriptomic analysis revealed significant upregulation of alpha-synuclein^9^ and downregulation of parkin^10^, which are critical to dopamine synthesis and degradation pathways, respectively. This distinct expression pattern suggests a disruption in the proteostasis network that could be pivotal in disease progression. Additionally, genes related to mitochondrial function showed decreased expression levels, consistent with the known impact of mitochondrial dysfunction in Parkinson’s pathology^11,12^. This differential gene expression or susceptibility aligns with the recent discovery of the distinct axonal projection patterns identified in dopaminergic neuron subtypes^6^. These patterns determine specific neural circuits affected during disease progression, providing a mechanistic basis for the diverse behavioral changes observed. For instance, subtypes projecting to motor-related areas may influence movement deficits, while those targeting limbic regions might underlie emotional or cognitive disturbances, illustrating how circuit-specific vulnerability translates into varied clinical manifestations^4–6^. However, it is not completely understood if specific molecularly identifiable subtypes of DANs in the ventral SNc are especially vulnerable in PD, and if the loss of function of these early vulnerable populations is just a biomarker of the disease or contributes to the early motor symptoms associated with PD.

Here, we characterized the progression of behavioral symptoms and dopaminergic degeneration in MitoPark mice, a well-established PD mouse model^13^, which replicates key aspects of the disease through the targeted deletion of the *Tfam* (mitochondrial transcription factor A) gene in DANs using the Cre-LoxP system. We characterized their motor behaviors in an open field with a motion sensor and identified bradykinesia and a tremor gradually emerging as MitoPark mice showed progressive dopaminergic degeneration in the SNc. We then performed single-nucleus RNA-sequencing (snRNA-seq) in MitoPark mice and matched littermate controls at different ages to identify transcriptomic changes in vulnerable dopaminergic populations, and correlated the susceptibility of different cell types to degeneration with the emergence of behavioral phenotypes. We identified specific dopaminergic cell types, highly ventrally localized *Sox6+* neurons, whose loss was timeline-wise correlated with the onset of motor symptoms. Further, we found that a subpopulation of the *Sox6+* neurons – the *Anxa1+* subset is more vulnerable than other *Sox6+* populations. We next created an Anxa1-Cre knock-in mouse to study the circuit architecture and function of this dopaminergic neuron subtype. We characterized the anatomical inputs and outputs and observed a specific pattern of strong projections of *Anxa1+* neurons to the dorsal striatum, not seen in other DAN populations, and observed that this subtype also receives different inputs than others. Finally, we showed that specific functional impairment of *Anxa1+* neurons is sufficient to induce slowness of movements and motor tremors, corresponding to the early PD-like symptoms. These results suggest that a specific vulnerable subtype of DANs is responsible for a range of early motor symptoms of PD, potentially highlighting the role of different dopaminergic neurons subtypes in the progression of the disease.

## Results

### Progressive development of PD-like motor deficits in MitoPark mice

The MitoPark mouse (genotype: *Tfam-loxP/loxP*, *+/DAT-cre*) is a transgenic model of Parkinson’s disease (PD) where the mitochondrial transcription factor A (*Tfam*) is removed selectively in dopaminergic neurons (DANs)^13^. Loss of *Tfam* in DANs leads to a progressive degeneration of these neurons and the manifestation of motor symptoms that are PD-like^13,14^.

We characterized spontaneous movement of MitoPark mice between 8 and 24 weeks of age in a square open field without any external cues or reward. These mice showed gradual development of locomotor deficits post-natal with decreased average speed (bradykinesia), increased immobility time and increased number of stops (akinesia) (**Fig. 1A-1D**). These effects were independent of environmental novelty, as similar differences were observed in both novel and habituated environments (**Fig. 1B-1D and Extended Data Fig. 1A and 1B**). We also performed a cylinder task to observe rearing behavior, which is more effortful and therefore more likely to be affected in PD, where there is a tendency towards less vigorous movements. We indeed confirmed a significant reduction in the number of rears in the older MitoPark mice (**Fig. 1E**). Given that typical PD has other motor symptoms, such as tremor in addition to bradykinesia and akinesia, we used a wireless motion sensor to detect subtle motor changes that are not easily observable by video-characterization. The sensor measures tri-axial acceleration with high resolution and allows for the continuous monitoring of movement with high frequency (200Hz)^15^, which could enable us to detect tremors that are usually poorly documented in genetically engineered rodent models of PD^16^. Given that tremor is defined as an involuntary rhythmic and oscillatory movement of a body part^17^, we explored spectral analysis of accelerometer data to quantify the potential emergence of abnormal oscillations with the progressive degeneration of DANs. Interestingly, we observed a specific oscillation within the 12-18 Hz frequency band that emerged in MitoPark mice at 16 weeks which gradually intensified with age (**Fig. 1F-1I and Extended Data Fig. 1C and 1D**). Notably, this oscillation was present even when the mice were immobile (**Fig. 1G and 1I**).

**Fig. 1.**
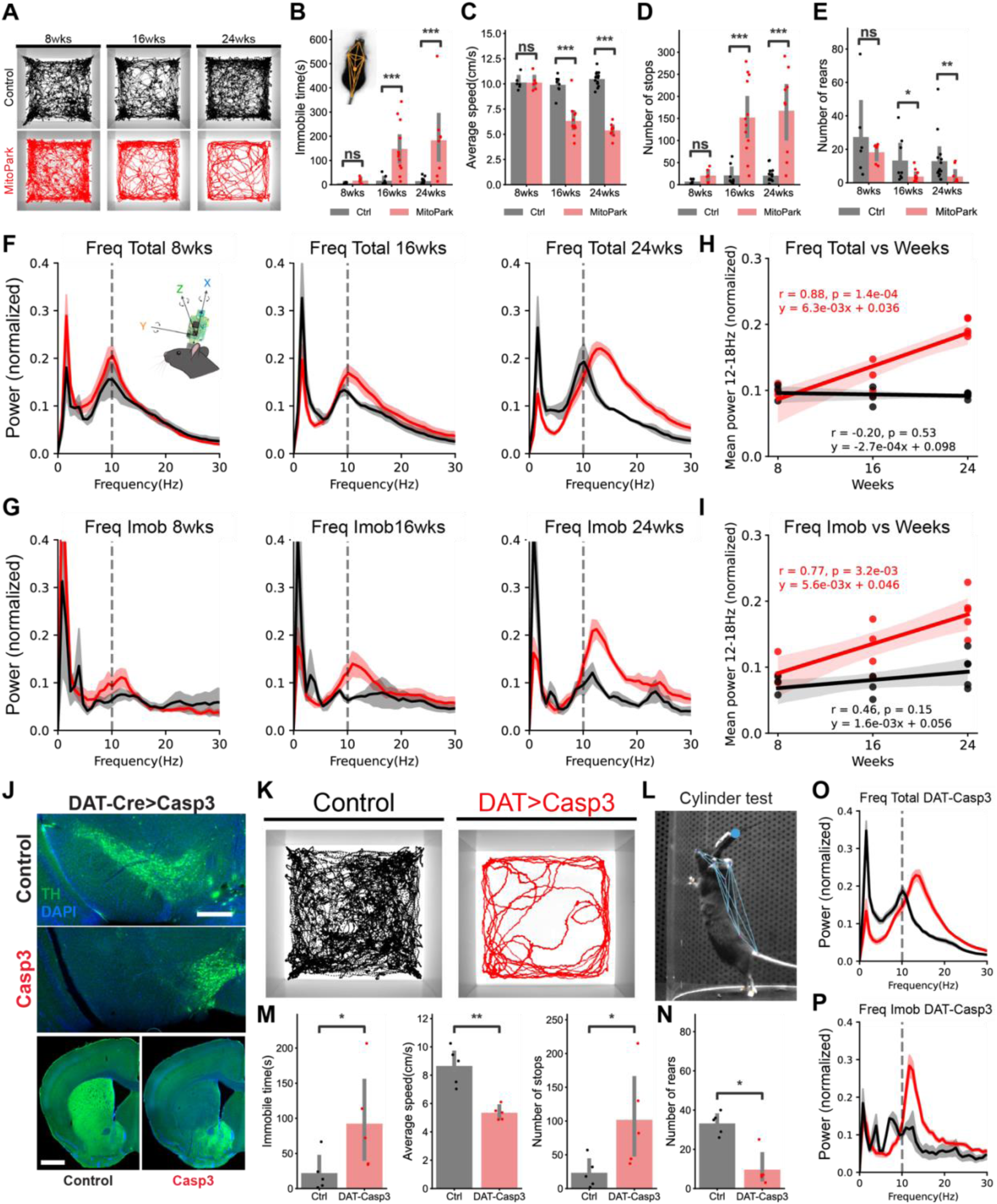
Progressive development of PD-like motor deficits in MitoPark mice. **A.** Cumulative trajectories of a representative MitoPark mouse and their control at different stages (8, 16 and 24wks) in the open-field test (black: control animals, red: MitoPark mice). **B-E**. Behavioral characterization of MitoPark mice and their littermate controls at each stage. **(B)** Immobility time: MitoPark mice at later stages spent significantly more time immobile compared to controls. The inset shows tracking performed using DeepLabCut. **(C)** Average speed during mobility: the speed of MitoPark mice was significantly lowered in 16 and 24wks. **(D)** Number of stops: the number of pauses in MitoPark mice showed significant differences compared to the controls. (**E**) Number of rears: MitoPark mice exhibited a significant reduction in rearing behavior at later stages. Statistical significance was determined by two-sided Mann–Whitney U-test (*p < 0.05, **p < 0.01, ***p < 0.001; ns = not statistically significant). Error bars show 95% confidence intervals. MitoPark 8weeks, n=6; Littermate controls 8weeks, n=6; MitoPark 16weeks, n=12; Littermate controls 16weeks, n=8; MitoPark 24weeks, n=9; Littermate controls 24weeks, n=15. **F-I.** Frequency analysis of total time **(F)** or immobility periods **(G)** at different ages of the animals. Mitopark mice showed a pronounced increase in the power of acceleration oscillations at the 12-18Hz range that becomes more prominent in the later stages. **(H, I)** The scatter plots show the correlation between age and mean power in the 12-18Hz frequency band for Mitopark mice and control animals during total period **(H)** or immobile period **(I)**. Pearson’s correlation coefficient reveals a strong positive correlation during both the total period (H: red, r=0.88, p<0.001) and the immobile period (I: red, r=0.77, p<0.01) in MitoPark mice. The solid line represents the linear regression model (H: red, y = 6.3e-03x + 0.036, R² = 0.78; I: red, y = 5.6e-03x + 0.036, R² = 0.60), demonstrating a statistically significant relationship between the two variables in MitoPark mice. MitoPark 8weeks, n=3; Littermate controls 8weeks, n=3; MitoPark 16weeks, n=4; Littermate controls 16weeks, n=3; MitoPark 24weeks, n=5; Littermate controls 24weeks, n=6. **J-P.** Injections of Casp3 virus in the SNc of DAT-Cre mice led to the similar phenotype of MitoPark mice 24 weeks. **(J)** The images demonstrated a specific reduction in dopamine neurons (DANs) within the SNc, as confirmed by staining with anti-TH antibody (green). **(K-N)** The DAT-Casp3 animals showed motor deficits characterized by diminished movement speed, increased incidences of halting and reduced the frequency of rearing behaviors. Statistical significance was determined by two-sided Mann–Whitney U-test (*p < 0.05; ns = not statistically significant). Error bars show 95% confidence intervals. DAT-Casp3 animals, n=5; Controls, n=5. **(O, P)** The virus injection led to the presentation of the same 12-18Hz oscillation seen in MitoPark mice at 24 weeks.

We next examined whether depletion of SNc DANs would be sufficient to lead to the development of PD-like symptoms similar to those observed in MitoPark mice. We therefore genetically ablated the neurons in the SNc of DAT-Cre mice by bilateral stereotactic injection of an adeno-associated virus (AAV) to express a Cre-dependent apoptotic gene Caspase-3 (AAV-Ef1a-FLEX-taCasp3)^18^ in the SNc of DAT-Cre mice. Selective losses of the SNc DANs and their projections were observed after 4 weeks of viral expression (**Fig. 1J**). We evaluated mice behavior in the open field, and found that the depletion of SNc DANs reproduced the PD-like phenotype observed in the 24 weeks of MitoPark mice (**Fig. 1A-1D, 1K, 1M and Extended Data Fig. 1E–1H**). We also observed a marked reduction in rearing behavior in the animals with ablation of SNc DANs (**Fig. 1L and 1N**). Furthermore, we also replicated the emergence of the tremor as observed in MitoPark mice (**Fig. 1O and 1P**). These results suggest that the neuronal loss in the SNc is sufficient to produce the bradykinesia, akinesia, and tremor observed in the MitoPark mice.

### Progressive loss of dopamine neurons in the ventral midbrain in MitoPark mice

To determine whether the mitochondrial *Tfam* deficit leads to changes in the number of midbrain DANs, we surveyed *Th* expression in MitoPark mice and monitored changes in *Th+* cells in the SNc of MitoPark mice at 8, 16 and 24 weeks (**Fig. 2A and 2B)**. We also examined *Th+* axonal projections from the SNc of MitoPark mice to striatum and performed brain reconstruction. Given that the current Allen Common Coordinate Framework (Allen CCF) does not have anatomical subdivisions of the striatum^19^, we created an updated map based on cortical projections^20^ and applied the imaging analysis pipeline BrainJ as previously described^21^ (**Extended Data Fig. 2**). This updated atlas allowed us to quantitatively analyze the density of projections neurons in subregions of the striatum and visualize the prominent loss of *Th+* axonal projections in MitoPark mice (**Fig. 2C and Extended Data Fig. 2C)**. We observed no observable differences in the projections from SNc DANs to the striatum at 8 weeks in MitoPark mice. However, by 16 weeks, their projections were significantly reduced compared to age-matched controls, especially for those targeting the dorsal striatum. By 24 weeks, the projections were largely depleted. (**Fig. 2A and 2C)**. Comparable progression has been reported in MCI (mitochondrial complex I)-Park mice in a previous study^12^.

**Fig. 2.**
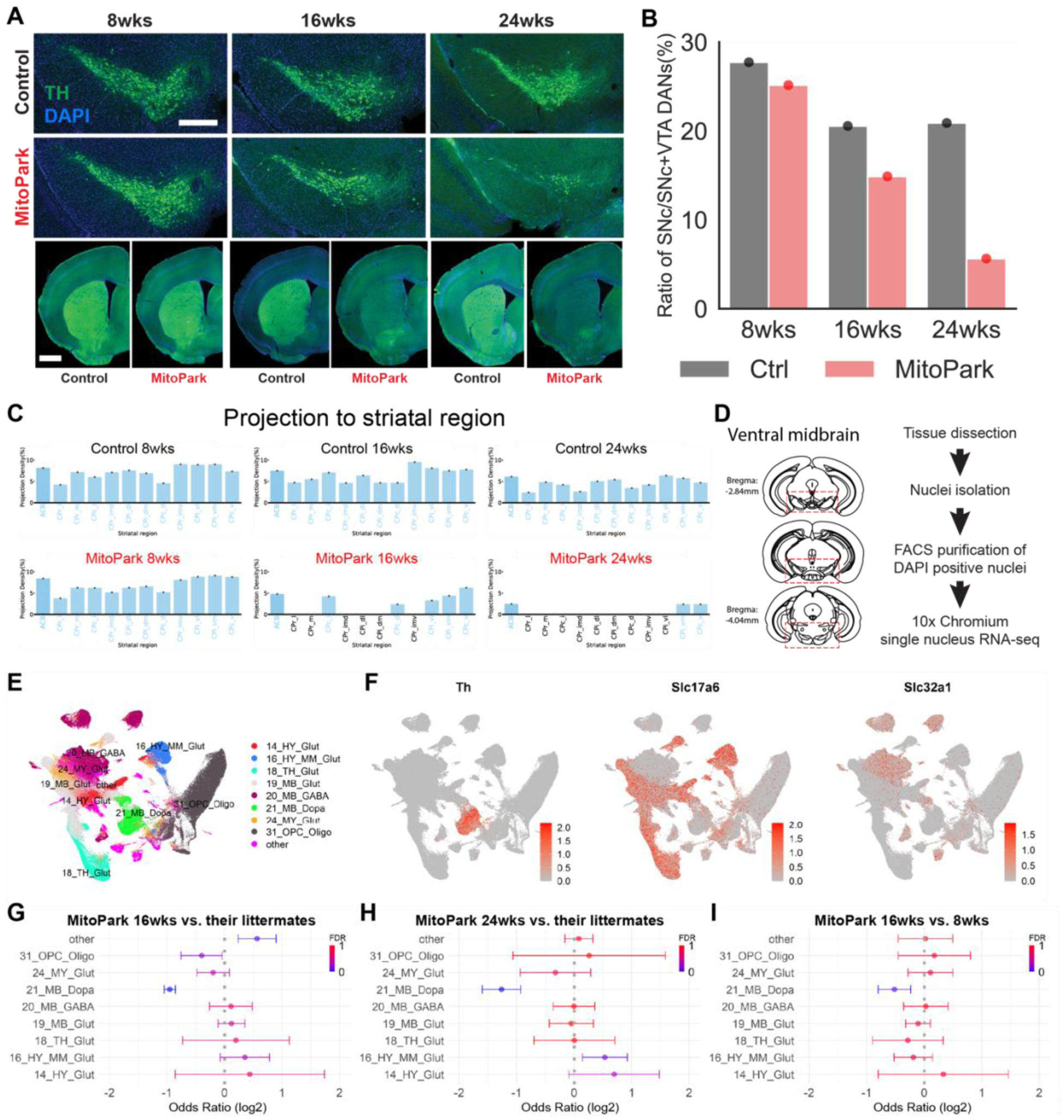
Progressive degeneration of dopamine neurons in MitoPark mice. **A-C.** TH immunohistochemistry of MitoPark mice and their littermate controls at 8, 16 and 24 weeks. (A) Histological images of the mice at each stage. Scale bars: 400μm for the substantia nigra (top) and 1mm for the striatum (bottom). (B-C) MitoPark mice exhibited a gradual and preferential loss of DANs in the substantia nigra, and their projections (each n=1). **D.** The workflow for single-nucleus RNA (snRNA) sequencing commenced with the collection of all tissues from the ventral midbrain (red dashed lines). The tissue samples underwent nuclei isolation, followed by the selection of DAPI+ nuclei. These nuclei were subsequently processed using the 10x Genomics platform for sequencing. **E.** UMAP visualization of snRNA-seq data from the ventral midbrain. Data are combined across all animals (MitoPark mice and their controls) and classified by the Allen Brain Cell (ABC) Atlas class annotations. Nuclei that are limited in number or belong to infrequent categories are designated as ‘others.’ **F.** Feature plot showing expression of marker genes used to label the main class of cells: *Th* (DANs), *Slc17a6* (excitatory neurons) and *Slc32a1* (inhibitory neurons). **G-I.** Dots and whiskers represent odds-ratio (OR) with 95% confidence interval (CI) obtained from MASC. OR estimates of major cell types associated with MitoPark mice (color; false discovery rate (FDR)-adjusted P<0.05). **(G)** MitoPark 16 weeks versus their littermate controls; 21_MB_Dopa (OR=-0.95, FDR-adjusted P<0.05). **(H)** MitoPark 24 weeks versus their littermate controls; 21_MB_Dopa (OR=-1.26, FDR-adjusted P<0.05). **(I)** MitoPark 16 weeks versus 8 weeks; 21_MB_Dopa (OR=-0.51, FDR-adjusted P=0.057).

We also employed single-nucleus RNA sequencing (snRNA-seq) of the ventral midbrain of MitoPark mice, their littermate controls (genotype: *Tfam-loxP/loxP*, *+/+* or *Tfam-loxP/+*, *+/+*) and C57BL/6 (B6) mice at 8, 16, and 24 weeks of age (8 mice in each group; 4 males and 4 females). We used a previously described experimental approach to isolate nuclei and profile their transcriptomes from mice^22^ (**Fig. 2D and Extended Data Fig. 3A**, see “Methods” section). We obtained a total of 293,435 nuclei, of which 93,703 were from MitoPark mice, 100,266 were from their littermate controls and 99,466 were from B6 mice. We mapped cell annotations to our data with the Allen Brain Cell (ABC) Atlas using their MapMyCells tool^23^ (**Fig. 2E and Extended Data Fig. 3B**). Dopaminergic nuclei were identified as the subclass annotation of 215_SNc_VTA_RAmb_Foxa1_Dopa at the ABC Atlas, within the class annotation of 21_MB_Dopa (**Extended Data Fig. 3B**). The dopaminergic population was also clearly distinguished by expression of the canonical marker gene, *Th* (**Fig. 2F**). Using *Rbfox3* to mark neuronal cells, we identified other neuronal cell types such as excitatory (*Slc17a6*) and inhibitory (*Slc32a1*) cell types (**Fig. 2F and Extended Data Fig. 3C**). Oligodendrocytes were characterized by the expression of *Mog*, while Oligodendrocyte precursor cells (OPCs) expressed high levels of *Pdgfra*. Microglia were characterized by *Ptprc* expression, while astrocytes and endothelial cells were marked by specific expression of *Gja1* and *Nostrin*, respectively (**Extended Data Fig. 3C**). Consistent with our histological findings, we observed a significant decrease in the number of DANs in MitoPark mice between 8 weeks and 24 weeks (**Extended Data Fig. 4A**). To address whether cell types are enriched or depleted in MitoPark mice, we compared the proportions of all cell-types across the ventral midbrain at each stage. We found that the dopaminergic populations significantly decreased with age compared with other cell types (**Extended Data Fig. 4B**). We also performed mixed-effects modeling (MASC)^24^ that accounts for technical and biological confounders to identify cellular populations associated with the disease models (**Fig. 2G-2I and Extended Data Fig. 4C-4E**). MitoPak mice had a significant reduction of DANs at 16 and 24 weeks relative to their littermate controls, while the other cell-types were stable (**Fig. 2G, 2H, Extended Data Fig. 4C and 4D**). A similar reduction of DANs was also observed between 8 and 16 weeks in the same genotype (**Fig. 2I and Extended Data Fig. 4E**). Together, these data demonstrate that *Tfam* depletion in MitoPark mice induces robust and selective dopaminergic degeneration without altering the number of other neighboring cell types (**Fig. 2G-2I and Extended Data Fig. 4B-4E**).

### Vulnerability of *Sox6+* population during the progression of PD-like phenotypes

We next aimed to identify changes within specific subtypes of DANs during the progression of neurodegeneration. We isolated 18,611 DANs from nuclei of the ventral midbrain, and snRNA-seq data was processed to identify distinct groups representing unique neuronal subtypes (**Fig. 3A, Extended Data Fig. 5A and 5B**). We identified 8 DAN subtypes and confirmed the expression of well-known subtype markers *Sox6*, *Slc17a6, and Calb1*^4^ (**Fig. 3B**). Each group was segregated by distinct genetic markers, which included known dopaminergic neuron subtype-specific genes (**Extended Data Fig. 5C and 6A-6C**). To determine which neuronal subtypes were most affected as neurodegeneration progressed, we analyzed the cell composition within each identified group from MitoPark mice and their age-matched controls. Comparative analysis showed significant changes in the composition of dopaminergic neuronal subtypes (**Fig. 3C**). Notably, the proportion of *Sox6+* neurons was markedly reduced in MitoPark mice, whereas the proportion of other neuron cell types such as *Megf11+* or *Synpr+* remained ‘stable’ throughout the degeneration process. As the overall number of dopamine neurons decreases in MitoPark mice, the proportion of these ‘stable’ neurons increased, suggesting they are more resilient compared to other dopamine subtypes that are more susceptible to degeneration (**Fig. 3C**). These neurons are mostly *Vglut2+* (*Slc17a6*) (**Fig. 3A and 3B**), which has previously been reported to play a potential neuroprotective role in the survival of DANs in PD model animals^25,26^. Additionally, there are dopamine neuron subtypes that undergo uniform degeneration over time (*Plekhg1+*, *Cck+*, *Sema5b+*, *Etv1+*, *Lypd1+*) (**Fig. 3C**). Using the MASC approach, we further analyzed the differential proportion of different cell-types or states across different groups (**Fig. 3D-3F**) while adjusting for age, sex, and batch as covariates. The *Sox6+* population showed a significant decrease in the odds ratio in MitoPark mice compared to their littermate controls, especially at 16 or 24 weeks (**Fig. 3D and 3E**), and it was not due to age-related effects alone (**Extended Data Fig. 6D-6G**). We also conducted the test across different ages within the same genotype to assess developmental changes in cell population dynamics (**Fig. 3F**). The odds ratio of *Sox6+* population still declined, suggesting that this subtype exhibits greater vulnerability than other DANs in PD, consistent with findings from human PD transcriptomic studies^8^.

**Fig. 3.**
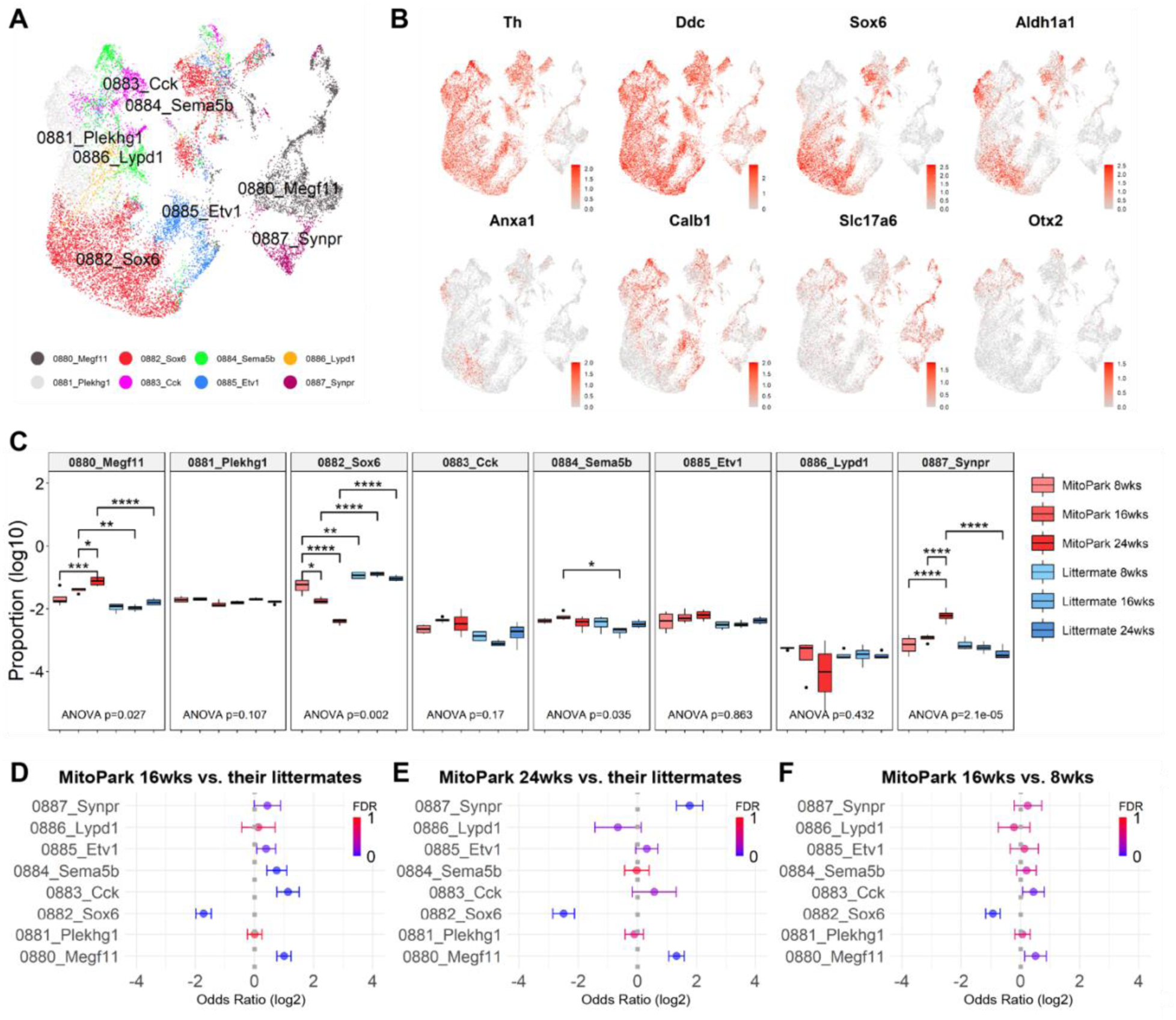
Subtype-specific degeneration within the dopaminergic populations. **A.** UMAP representation of 18,611 dopamine neuron nuclei, colored by the supertype annotation of the Allen Brain Cell (ABC) Atlas. In supertype annotation of the ABC Atlas, DANs are largely classified into 8 types. **B.** Representative feature plots showing gene expression associated with dopamine used to label the main class of cells. **C.** Distribution of dopamine subtype proportions across MitoPark and their control animals. 0882_Sox6 is mostly affected among other dopamine neuron subtypes. Statistical significance was determined by two-way ANOVA (p-value indicates the interaction effect between genotype and age) followed by Tukey’s HSD test for multiple comparisons (*p < 0.05, **p < 0.01, ***p < 0.001, ****p < 0.0001). **D-F.** Odds-ratio estimates of each dopamine subpopulation in **(D)** MitoPark 16 weeks versus their littermate controls; 0882_Sox6 (OR=-1.72, FDR-adjusted P<0.05), 0880_Megf11 (OR=1.00, FDR-adjusted P<0.05) and 0883_Cck (OR=1.13, FDR-adjusted P<0.05). **(E)** MitoPark 24 weeks versus their littermate controls; 0882_Sox6 (OR=-2.49, FDR-adjusted P<0.05), 0887_Synpr (OR=1.76, FDR-adjusted P<0.05) and 0880_Megf11 (OR=1.31, FDR-adjusted P<0.05). **(F)** MitoPark 16 weeks versus 8 weeks; 0882_Sox6 (OR=-0.93, FDR-adjusted P=0.057).

### A subset of *Sox6+* DANs are more vulnerable in MitoPark mice

*Sox6* is involved in the development and differentiation of DANs, particularly in the ventral midbrain DANs^27–29^. The *Sox6+* supertype (ID: 0882 in 215_SNc_VTA_RAmb_Foxa1_Dopa) can be further subdivided according to their anatomical location, distinct molecular signatures, and functional properties^4^. To explore the diversity of *Sox6+* neurons, we conducted transcriptomic analysis of the subtypes and performed the cluster annotation based on the ABC atlas^23^. We found 43 clusters (ID: 3837-3879) under the supertype (ID: 0880-0887) annotation, of which, the *Sox6+* supertype can be divided into 7 subsets (ID: 3853-3859) (**Extended Data Fig. 5A**). We quantified the proportions of *Sox6+* dopaminergic subtypes in MitoPark and control animals. Two subsets, *Anxa1+* (ID: 3857) and *Arpp21+* (ID: 3859), showed statistically significant reductions compared to controls (two-way ANOVA followed by Tukey’s HSD). *Anxa1+* was significantly reduced across age, whereas *Arpp21+* reached significance only at 24 weeks (**Fig. 4A**). Although other *Sox6+* subsets showed a tendency toward reduced proportions in MitoPark animals, this pattern prompted the use of a complementary differential abundance analysis. We therefore performed an additional differential abundance analysis using Milo, a method that offers a structured approach to exploring variations in cell compositions across diverse experimental conditions^30^ (**Fig. 4B-4J and Extended Data Fig. 7A-7I**). Milo analysis confirmed a significant depletion of *Sox6+* population in MitoPark mice at 16 and 24 weeks compared to their littermate controls, consistent with previous analyses (**Fig. 3C-3F and Extended Data Fig. 7A-7I**). Following cluster annotation, most *Sox6+* subsets exhibited pronounced reductions in gene expression at 16 and 24 weeks (log fold change < −3 at 16 weeks and < −5 at 24 weeks; **Fig. 4E, 4F, 4H and 4I**). Additionally, comparison between 8 and 16 weeks in MitoPark animals revealed two subsets (*Grin2c+* (ID: 3854) and *Anxa1+* (ID: 3857)) with robust gene expression decreases (log fold change < −2; **Fig. 4D, 4G, 4J, and Extended Data Fig. 7J-7L**). An additional analysis using a linear mixed-effects model (LMM)^31^, followed by Tukey’s post hoc comparisons, showed that the 3857_Anxa1 cluster consistently exhibited the strongest and most statistically significant age-by-genotype interaction, and further indicated that this cluster is already slightly affected at 8 weeks (MitoPark vs. Control; 8 weeks: 2.37E-03, 16 weeks: 3.34E-06, 24 weeks: 1.41E-04)^31^. We also compared DAT expression across *Sox6+* dopaminergic subtypes and found no significant differences, indicating that selective vulnerability is unlikely to be driven by DAT levels (**Extended Data Fig. 7M**). To verify the transcriptional reduction of DAN subsets in the SNc of MitoPark mice, we conducted immunostaining with an antibody targeting *Aldh1a1*, a well-established marker for ventral DANs^4,5^ (**Fig. 4K**). *Aldh1a1+* neurons constitute a relatively large neuronal subset and are recognized as selectively vulnerable in PD, encompassing the *Anxa1+* population in the ventral SNc^5,32–34^. We found a remarkable loss of *Aldh1a1*+ DANs in MitoPark mice at 16 weeks, particularly in the rostral-ventral region (**Fig. 4K**). These results demonstrate that *Sox6+* DANs exhibited differential susceptibility, with *Anxa1+* DANs emerging as one of the earliest populations to show vulnerability relative to other DANs.

**Fig. 4.**
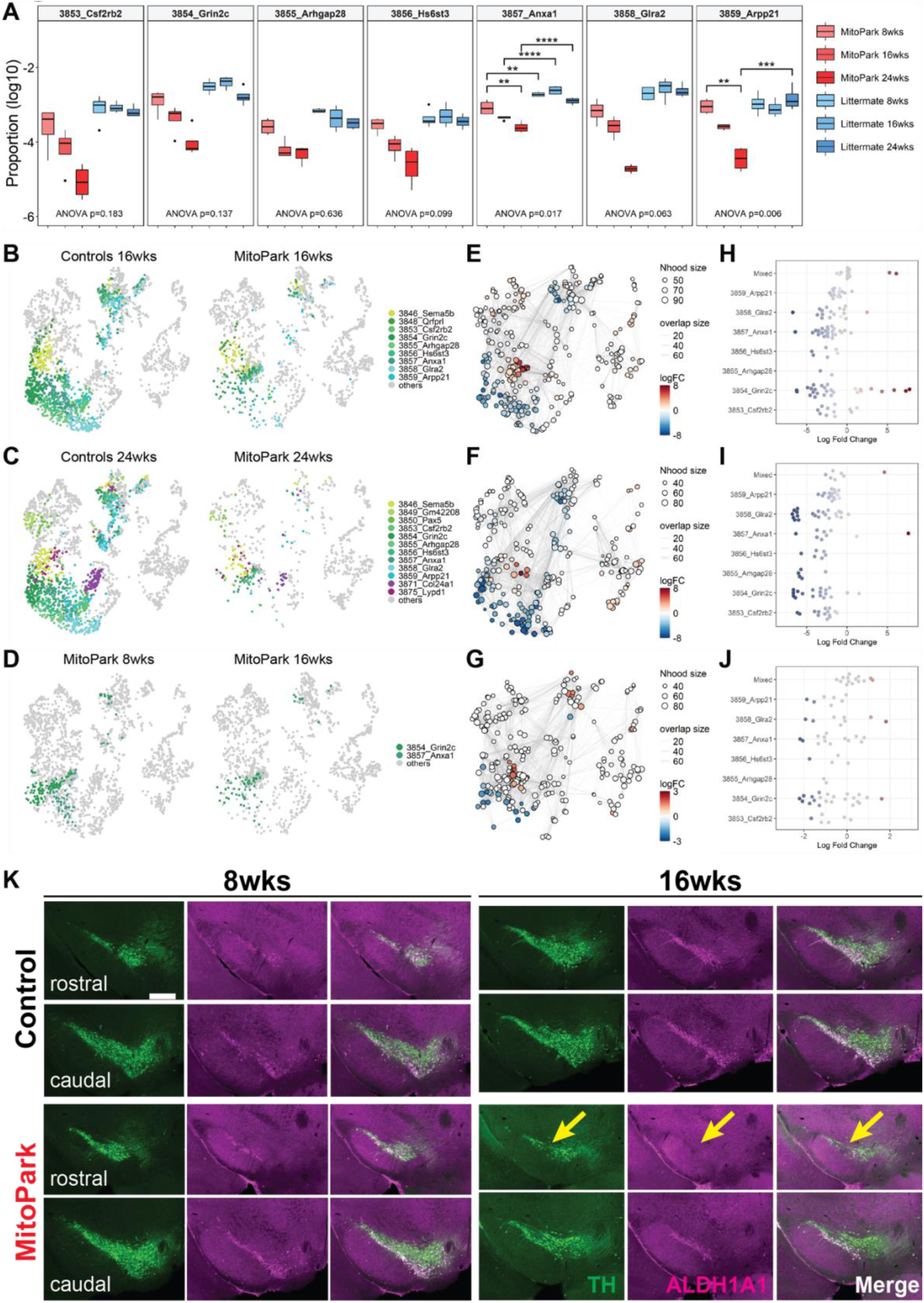
A subset of *Sox6+* population is more vulnerable than others in MitoPark mice. Differential abundance analysis for the dopamine clusters of MitoPark mice and their littermate controls. The analysis reveals a subtype-specific reduction in MitoPark mice at a later stage. **A.** Distribution of *Sox6+* dopamine subtype proportions across MitoPark and their control animals. 3857_Anxa1 is significantly affected among other dopamine neuron subtypes at 16 and 24 weeks. Statistical significance was determined by two-way ANOVA (p-value indicates the interaction effect between genotype and age) followed by Tukey’s HSD test for multiple comparisons (*p < 0.05, **p < 0.01, ***p < 0.001, ****p < 0.0001). **B-D.** UMAP plots illustrating dopamine neuron subpopulations in MitoPark mice and their littermate controls (n = 8 per genotype and stage), highlighting clusters with significant differential abundance (log fold change < 2). The gene groups affected across all comparisons are shown: (B) Controls vs. MitoPark at 16 weeks, (C) Controls vs. MitoPark at 24 weeks, and (D) MitoPark at 8 vs. 16 weeks, as identified by the differential abundance analysis. **E-G.** Neighborhood (Nhood) graphs with the results from Milo differential abundance testing between MitoPark mice and controls (E:16 weeks, F: 24 weeks), and between MitoPark mice at different stages (G: 8 and 16 weeks). Nodes represent neighborhoods, colored by their log fold change compared to their controls (blue: less abundant, red: more abundant, gray: non-differentially abundant). Graph edges show the number of cells shared between two neighborhoods. **H-J**. Beeswarm plots display the distribution of log-fold changes in cell abundance. Dots representing neighborhoods that overlap with the same cell populations are grouped together. Colors are represented similarly to **E-G**. **K.** Histological images representing 8 and 16 weeks of MitoPark mice reveal that significant cell depletion in the ventral region, particularly in the rostral part of the SNc (TH: green, ALDH1A1: magenta). Scale bar 400μm.

### Whole-brain efferent and afferent connectivity of *Anxa1+* DANs

The results above indicate a subset of *Sox6+* DANs, the *Anxa1+* neurons, is more vulnerable than others. However, it remains unclear whether loss of these neurons is an early biomarker of disease, or are actually driving early symptoms in PD. To address this, we generated an Anxa1-Cre knock-in mouse line (see ‘Method’ section), with Cre recombinase under transcriptional control of an Anxa1-specific promoter and characterized its expression using GFP to visualize cell bodies and axon projections to output structures. To achieve this, we injected Cre-dependent AAV-GFP into the SNc of Anxa1-Cre mice and counted the number of *Th+* cells with *Calb1* antibody, which labels the dorsal dopaminergic populations in the SNc (**Fig. 5A and 5B**). In all mice, we observed robust co-localization of GFP with *Th+* cells but not *Calb1+* cells, suggesting that most of the GFP+ cells in the SNc are ventral specific DANs (**Fig. 5A and 5B**). We then examined GFP+ axonal projections from the SNc of all DANs or *Anxa1+* DANs and performed whole brain reconstruction using BrainJ (**Fig. 5C and 5D)**. The quantitative analysis revealed prominent axonal projections of *Anxa1+* DANs to the dorsal striatum (**Fig. 5D**). We also created a DAT-Flp knock-in mouse line^35^ (**Extended Data Fig. 9A;** see ‘Method’ section) and compared the axonal projections of SNc *Anxa1+* DANs with other DANs subtypes such as *Vglut2+* or *Calb1+* (**Extended Data Fig. 9B**). As shown previously^6^, these subsets have distinct axonal projections to the striatum; *Vglut2+* DANs project to the dorsolateral and tail striatum, while *Calb1+* DANs project to the ventral striatum (**Extended Data Fig. 9B**). We also quantitatively analyzed SNc *Vglut2+* and *Calb1+* DAN axonal projections throughout the whole brain, which make specific projections to many different regions. The fact that different populations of DANs have such distinctive projections opens the possibility that different populations modulate different behaviors or behavioral parameters^5^ (**Extended Data Fig. 9C)**.

**Fig. 5.**
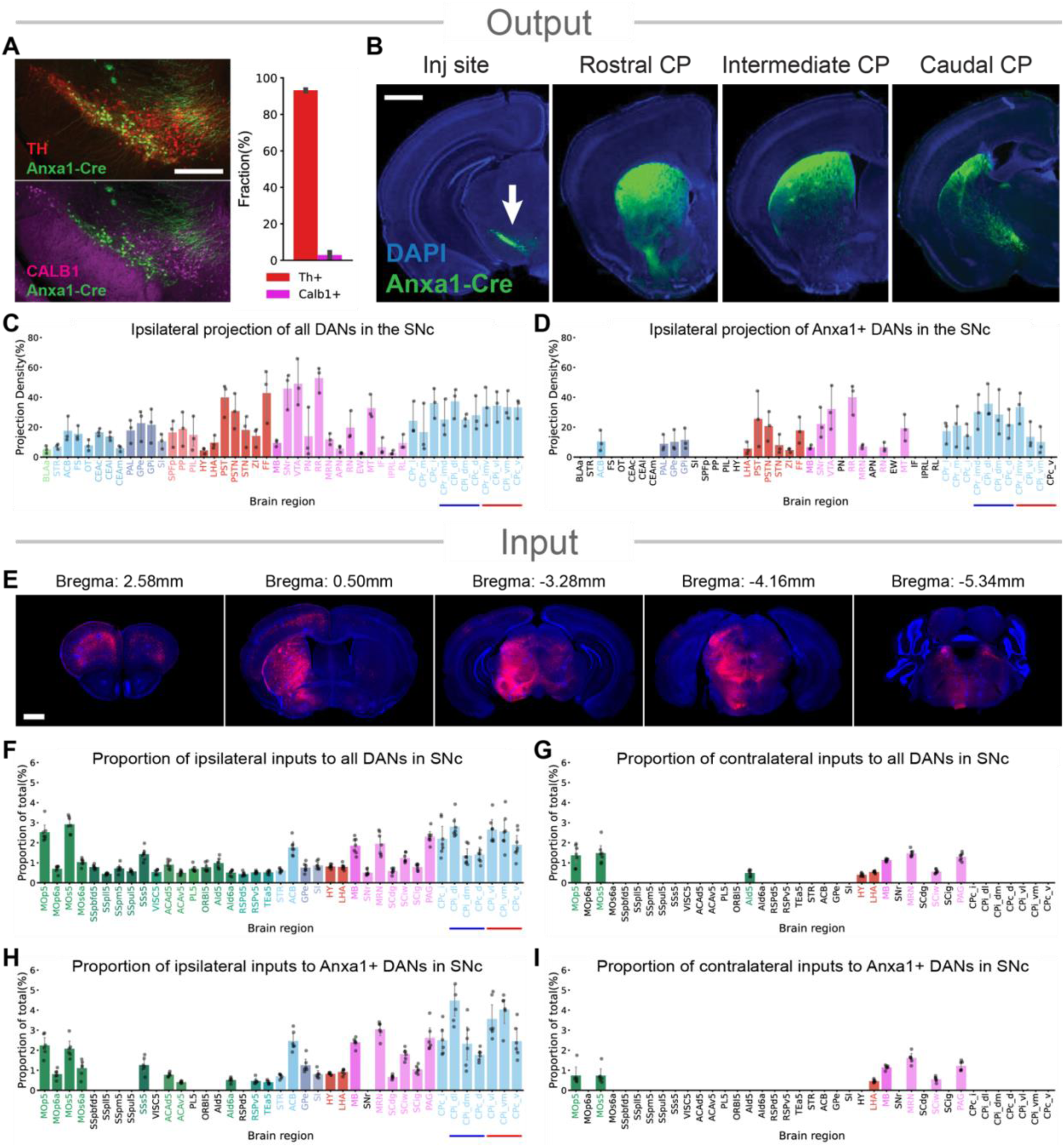
Input-output circuit architecture of *Anxa1+* DANs in the SNc. **A.** In the Anxa1-Cre line, the majority of neurons within the SNc are positive for *Th* and negative for *Calbindin1* (TH, red; CALB1, magenta; n=2). Scale bars: 400μm. **B.** Axonal projections of *Anxa1+* DANs in the SNc. They preferentially project to dorsal striatum that integrates sensorimotor information from cortical and thalamic regions. Scale bars: 1mm. **C-D.** Brain regions to which all DANs or *Anxa1+* DANs in the SNc project, measured as the fraction of neurites found within those brain structures defined Allen brain atlas (each n=3). Blue horizontal line indicates dorsal striatum and the red line indicates ventral striatum. **E.** Representative images of rabies infected cells (tdTomato). Rabies virus was injected in the SNc of Anxa1-Cre counterstained with DAPI in blue. Scale bar 1mm. **F-I.** Statistical analysis of the whole-brain distribution of ipsilateral (left) or contralateral (right) monosynaptic inputs to all DANs (F, G) or *Anxa1+* dopamine neuron subtype (H, I) in the SNc. Average proportion of tdTomato-labeled neurons in approximately 50 brain regions, each with more than 200 cells, was greater than 0.2% of the total input to DANs in DAT-Cre mice (n=7) and Anxa1-Cre mice (n=5; 4 animals for Anxa1-Cre+ and 1 animal for DAT-Flp+;Anxa1-Cre+). Brain areas are color-coded by the Allen Brain Atlas. **[Acronym]** ACAd5, Anterior cingulate area, dorsal part, layer 5; ACAv5, Anterior cingulate area, ventral part, layer 5; ACB, Nucleus accumbens; AId5, Agranular insular area, dorsal part, layer 5; AId6a, Agranular insular area, dorsal part, layer 6a; APN, Anterior pretectal nucleus; BLAa, Basolateral amygdalar nucleus, anterior part; CEAc, Central amygdalar nucleus, capsular part; CEAl, Central amygdalar nucleus, lateral part; CEAm, Central amygdalar nucleus, medial part; CPc_d, Caudal Caudoputamen, dorsal; CPc_i, Caudal Caudoputamen, intermediate; CPc_v, Caudal Caudoputamen, ventral; CPi_dl, Intermediate Caudoputamen, dorsolateral; CPi_dm, Intermediate Caudoputamen, dorsomedial; CPi_vl, Intermediate Caudoputamen, ventrolateral; CPi_vm, Intermediate Caudoputamen, ventromedial; CPr_imd, Rostral Caudoputamen, intermediate dorsal; CPr_imv, Rostral Caudoputamen, intermediate ventral; CPr_l, Rostral Caudoputamen, lateral; CPr_m, Rostral Caudoputamen, medial; EW, EdingerWestphal nucleus; FF, Fields of Forel; FS, Fundus of striatum; Gpe, Globus pallidus, external segment; Gpi, Globus pallidus, internal segment; HY, Hypothalamus; IF, Interfascicular nucleus raphe; IPRL, Interpeduncular nucleus, rostrolateral; LHA, Lateral hypothalamic area; MB, Midbrain; MOp5, Primary motor area, Layer 5; MOp6a, Primary motor area, Layer 6a; MOs5, Secondary motor area, layer 5; MOs6a, Secondary motor area, layer 6a; MRN, Midbrain reticular nucleus; MT, Medial terminal nucleus of the accessory optic tract; ORBl5, Orbital area, lateral part, layer 5; OT, Olfactory tubercle; PAG, Periaqueductal gray; PAL, Pallidum; PIL, Posterior intralaminar thalamic nucleus; PL5, Prelimbic area, layer 5; PN, Paranigral nucleus; PP, Peripeduncular nucleus; PST, Preparasubthalamic nucleus; PSTN, Parasubthalamic nucleus; RL, Rostral linear nucleus raphe; RN, Red nucleus; RR, Midbrain reticular nucleus, retrorubral area; RSPd5, Retrosplenial area, dorsal part, layer 5; RSPv5, Retrosplenial area, ventral part, layer 5; SCdg, Superior colliculus, motor related, deep gray layer; Scig, Superior colliculus, motor related, intermediate gray layer; Sciw, Superior colliculus, motor related, intermediate white layer; SI, Substantia innominata; SNr, Substantia nigra, reticular part; SPFp, Subparafascicular nucleus, parvicellular part; SSpbfd5, Primary somatosensory area, barrel field, layer 5; SSpll5, Primary somatosensory area, lower limb, layer 5; SSpm5, Primary somatosensory area, mouth, layer 5; SSpul5, Primary somatosensory area, upper limb, layer 5; SSs5, Supplemental somatosensory area, layer 5; STN, Subthalamic nucleus; STR, Striatum; TEa5, Temporal association areas, layer 5; VISC5, Visceral area, layer 5; VTA, Ventral tegmental area; ZI, Zona incerta

To identify the afferent inputs onto *Anxa1+* dopaminergic neuronal subtypes and how these inputs differ from other DAN subtypes, we used CVS-N2c rabies tracing, a strain with enhanced transsynaptic labeling^36^, with DAT-Cre+, Anxa1-Cre+ (and also DAT-Flp+;Anxa1-Cre+), DAT-Flp+;Vglut2-Cre+ and DAT-Flp+;Calb1-Cre+ mice. We first examined the inputs to all DANs in the SNc by injecting two Cre-dependent helper viruses (TVA and N2cG) to express the avian receptor (TVA protein) and the rabies glycoprotein G (N2cG) unilaterally into the SNc of DAT-Cre+ mice. Two weeks later, CVS rabies virus (CVS-tdTomato) was injected into the same site (**Extended Data Fig. 10A;** see “Methods” section). We observed tdTomato-labeled neurons in more brain regions than previous studies^37,38^, especially detected much more inputs for cortical regions, likely attributed to the higher transsynaptic property of the N2C strain. These labeled neurons, primarily restricted ipsilateral to the injection site, represent monosynaptic afferent inputs to the SNc DANs (**Fig. 5F, 5G and Extended Data Fig. 10B, 10C**). The same viral injection strategy was also used using Anxa1-Cre+ (and DAT-Flp+;Anxa1-Cre+), DAT-Flp+;Vglut2-Cre+, and DAT-Flp+;Calb1-Cre+ mice, which also labeled an abundant number of tdTomato+ neurons in various parts of brain regions (**Fig. 5E, 5H, 5I and Extended Data Fig. 9D-9G**). The dopaminergic neuron subtypes exhibit modest differences in their ipsilateral inputs, particularly with respect to inputs from cortical regions. For example, while *Anxa1+* DANs receive strong inputs from the motor cortex, this subtype does not receive significant inputs from the somatosensory, prelimbic, or orbital cortex. In contrast, *Vglut2+* DANs receive minimal inputs from the ventral regions of the anterior cingulate and retrosplenial cortices compared to other dopamine subtypes. In most other cases, however, *Anxa1+* DANs receive similar inputs to the other dopaminergic subtypes, suggesting a far more shared organization of inputs into SNc dopaminergic circuits than their outputs (**Fig. 5E-5I and Extended Data Fig. 9D-9G**).

### Inhibition of *Anxa1+* DANs is sufficient to cause early PD-like motor deficits

The activity of *Anxa1+* neurons has been shown to correlate with movement acceleration^5^. To determine whether the loss of *Anxa1+* neurons represents an early biomarker of disease or contributes causally to early PD symptoms, we selectively silenced synaptic release from *Anxa1+* dopaminergic neurons using an AAV expressing tetanus toxin^39^ (AAV-FLEX-TeLC-EYFP; **Fig. 6A**). We found that silencing of *Anxa1+* neurons caused a reduction in locomotor speed, suggesting it is sufficient to cause bradykinesia. However, it did not affect immobility time or number of stops, suggesting that it was not sufficient to cause akinesia. (**Fig. 6B-6D**). In a cylinder task, we observed the significant drops in the number of rears, suggesting that *Anxa1+* dopaminergic neuron activity is important for effortful movements (**Fig. 6C)**. We then performed spectral analysis on mice depleted of *Anxa1+* SNc dopaminergic neurons and found a slight shift in the frequency of tremor, resembling the phenotype of MitoPark mice at 16 weeks (**Fig. 6E**).

**Fig. 6.**
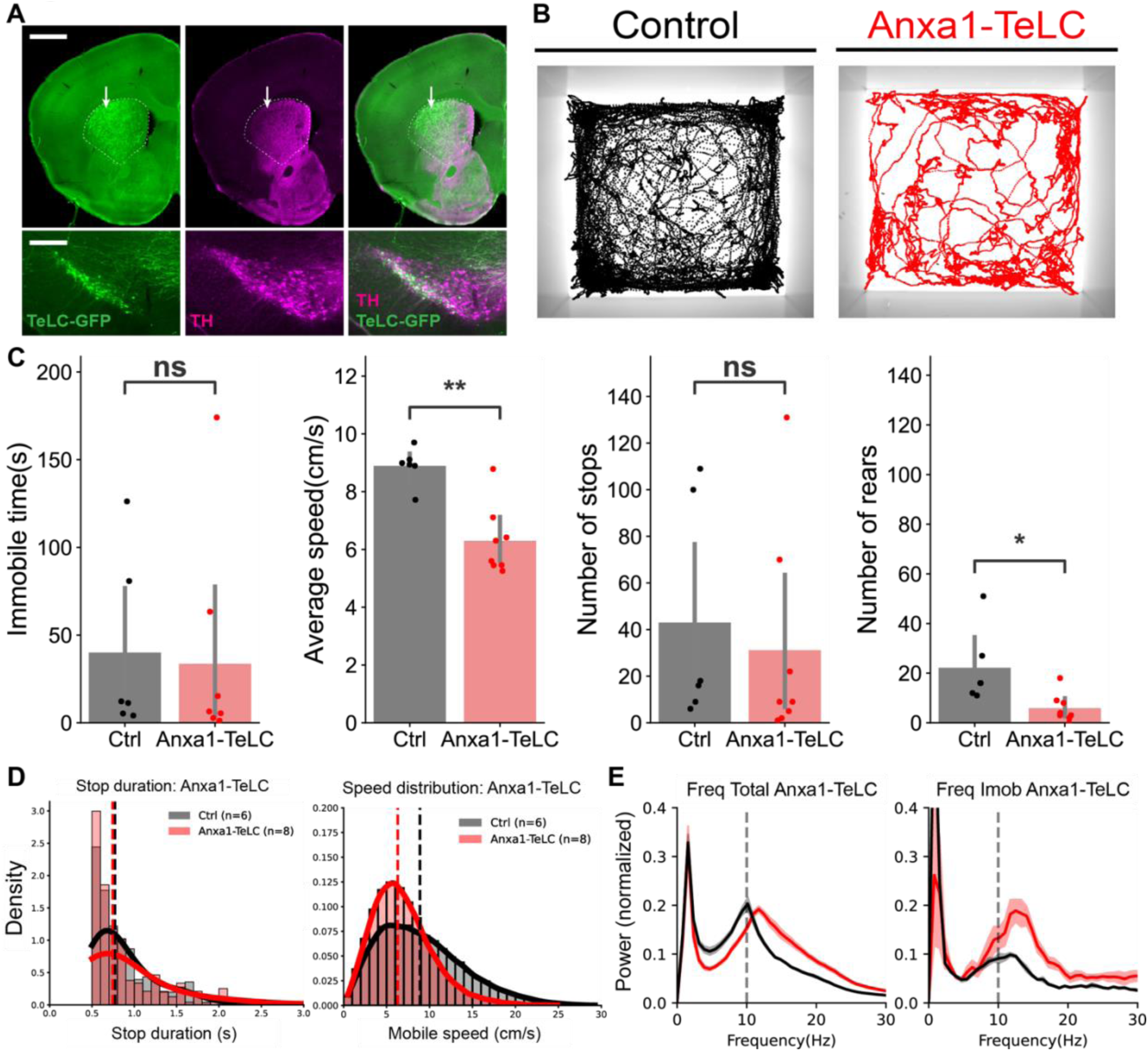
Inhibition of *Anxa1+* DANs leads to motor deficits resembling early onset PD-like symptoms. **A.** In the Anxa1-Cre line, targeted expression of the tetanus toxin light chain (TeLC) specifically eliminated dopamine synaptic transmission in the dorsal striatum (white arrows). Scale bars: 1mm for the striatum (top) and 400μm for the substantia nigra (bottom). **B-D.** Behavioral characterization of Anxa1-TeLC animals. (B) Cumulative trajectories of a representative Anxa1-TeLC mouse and their control in the open-field test (black: a control animal, red: Anxa1-TeLC mouse). (C) Anxa1-TeLC animals demonstrated slowed movement; however, there were no significant differences observed in terms of immobility duration or the frequency of stops. The frequency of rearing behaviors in the Anxa1-TeLC animals was significantly reduced. Statistical significance was determined by two-sided Mann–Whitney U-test (**p < 0.01, *p < 0.05; ns = not statistically significant). Error bars show 95% confidence intervals. Anxa1-TeLC animals, n=5; Controls, n=4. (D) This figure shows the probability density function (PDF) for stop duration (left) and speed distribution (right), with duration (in seconds) or speed (cm/s) on the x-axis and probability density on the y-axis. There is no significant difference in the distribution of stop durations between Anxa1-TeLC and control animals, although the speed distribution in Anxa1-TeLC animals is slightly shifted to the left, indicating slower movement. **E.** Anxa1-TeLC animals showed the similar 12-18Hz oscillation seen in MitoPark mice at 16 weeks.

Learning is also known to be affected early in PD and in MitoPark animals^40,41^. Given the early vulnerability of *Anxa1+* neurons, we wanted to determine whether their loss is sufficient to cause learning deficits, or if other neuronal populations are responsible. We used a Pavlovian conditioning task in which mice learned to lick for a reward when presented with a light cue. In this paradigm, a visual cue (CS+; LED stimulus) signaled the availability of a sucrose reward, delivered at random intervals, whereas an auditory cue (CS–; sound) was never paired with reward (**Extended Data Fig. 11A**). Thus, CS+ presentations should elicit increased reward expectation, reflected by elevated licking despite the unpredictability of cue timing. Across conditioning days, control mice showed a progressive increase in licking following CS+ presentation, indicating successful learning of the cue–reward association. In contrast, Anxa1-TeLC mice displayed markedly reduced licking responses (**Extended Data Fig. 11B**). To determine whether these effects were due to impairment in licking, we also analyzed poke events (**Extended Data Fig. 11C**). Control mice showed prompt, cue-triggered poking beginning on Day 2, whereas Anxa1-TeLC mice exhibited random, non–cue-directed poking. Total lick counts across each session did not differ between groups, and total poke counts showed a similar pattern except on Day 1 (**Extended Data Fig. 11D and 11F**). However, both the first-lick and first-poke latencies were significantly prolonged in Anxa1-TeLC mice, although these delays gradually decreased over days (**Extended Data Fig. 11E and 11G**).

Taken together, these data show that inhibition of *Anxa1+* DANs is sufficient to induce early PD-like features such as reduced movement speed, diminished movement vigor, tremor, and learning deficits, yet not sufficient to produce akinesia.

## Discussion

The data presented in this study revealed that a subtype of dopaminergic neurons (DANs) is selectively vulnerable in a mouse model of Parkinson’s disease (PD) and that its loss is sufficient to drive PD-like motor symptoms that are observed early in the disease. We conducted a detailed longitudinal analysis of the behavioral changes exhibited by MitoPark mice, a well-established PD model^13^. We identified progressive changes in bradykinesia, akinesia, and tremor. Concomitantly, these mice demonstrated a progressive degeneration of DANs in the midbrain, specifically within the substantia nigra pars compacta (SNc). These phenotypes observed in late-stage MitoPark mice were replicated by ablating DANs in the SNc. Additionally, we performed a comprehensive characterization of the molecular profiles of midbrain neurons as the disease progressed, and uncovered selective vulnerability in the *Sox6+* DANs, and in particular in the *Anxa1+* subtype. Finally, we observed that silencing this dopamine subtype was sufficient to produce bradykinesia and tremor of the magnitude observed early in the progression of the degeneration, but not akinesia. This reveals that the loss of these vulnerable neurons is not only a biomarker of early disease, it also drives the early motor symptoms observed early in the disease.

Our behavioral analysis rigorously mapped the progression of motor symptoms in MitoPark mice revealing early symptoms that were not previously reported. Furthermore, it revealed a previous unobserved phenotype associated with dopamine loss. By using a motion sensor in an open field behavioral assay, we have enhanced our capability to detect more subtle motor phenotypes that are not detected using conventional video-based characterization methods. This approach led us to identify tremors in MitoPark mice, which developed incrementally with the progressive degeneration of DANs in the brain. Furthermore, specific ablation and silencing of SNc DANs and the *Anxa1+* subtype revealed their important role in this tremor phenotype. Although the significance of the oscillation frequency we observed remains under investigation, the ∼10 Hz signal in control mice most likely reflects a physiological tremor linked to postural stabilization and the natural resonance of the sensorimotor system. In contrast, the tremor seen in MitoPark and dopamine-lesioned mice even during rest suggests the emergence of a central, PD-related tremor or increased rigidity that shifts the mechanical tremor frequency. Notably, the tremor frequencies in MitoPark mice do not fall within the typical 4-6 Hz range observed in PD patients^42^; however, species-specific differences in tremor frequency are well documented, as illustrated by the harmaline model of essential tremor, which produces oscillations of 8–10 Hz in monkeys, 10–12 Hz in rats, and 11–14 Hz in mice^43,44^.

Considering that there are several types of tremors observed in humans and the extent to which these tremors resemble those seen in human diseases has not yet been fully established, a more comprehensive characterization is imperative. Another key discrepancy between human PD and PD mouse models is that, in early-stage patients, motor symptoms such as resting tremor and bradykinesia typically emerge asymmetrically due to greater dopaminergic neuron loss in one hemisphere, producing a unilateral and focal onset that later becomes bilateral^45^. In contrast, PD mouse models generally exhibit more symmetric dopaminergic degeneration, resulting in bilateral, whole-body tremors and limiting the detection of focal symptom onset. The human and mouse musculoskeletal systems also differ substantially, which may contribute to the distinct tremor frequencies observed and reflect species-specific postural demands. This interpretation is supported by a recent study that employed similar inertial sensors in patients with PD^46^. These differences likely arise from both species-specific pathological features and the whole-body nature of our behavioral assessments.

In previous studies, *Aldh1a1+* DANs have been shown to be selectively vulnerable in PD^32^. Their properties including connectivity and firing patterns have been extensively characterized, and these works have highlighted their relevance to motor control and PD pathophysiology^32–34^. It has also been shown that *Aldh1a1+* neurons comprise distinct genetic and functional subsets^5^, which motivated us to perform independent molecular profiling. We performed an extensive longitudinal molecular profiling of the cell types in the ventral midbrain as the symptoms progressed in MitoPark mice, utilizing a newly developed atlas, the Allen Brain Cell (ABC) Atlas, for annotation^23^. This detailed examination identified a vulnerable population of DANs, *Sox6+*, the characteristics of which align closely with those identified in previous human and mouse studies^8^. In contrast, other subtypes, such as *Megf11+* (ID: 0880), and *Synpr+* (ID: 0887) displayed resilience to degeneration. We identified several additional subtypes of *Sox6+* DANs, with different vulnerability. *Anxa1+* (ID: 3857) neurons showed the earliest significant decrease compared to the littermate controls, but *Arpp21+* (ID: 3859) also showed vulnerability in late stages compared to other subtypes of *Sox6+* neurons. This early vulnerability was also observed in other mouse models and in humans in an accompanying study^47–49^. This variance in vulnerability and resilience indicates that the degeneration of DANs occurs at different rates in different types, and that the molecular composition of these subtypes may underly the mechanisms for vulnerability and resilience. Consequently, a deeper understanding of the molecular processes driving the degeneration and resilience of these various DAN subtypes is critical for understanding the progression of the disease, and could be harnessed for the development of biomarkers to monitor progression and eventually the response to disease modifying therapies.

We also characterized with unprecedented depth the afferent and efferent connectivity of three distinct subtypes of DANs in the SNc, roughly representing lateral, dorsal, and ventral SNc DANs, identified by the expression of genes *Vglut2+*, *Calb1+*, and *Anxa1+* respectively. Consistent with previous reports, these subtypes display distinct projection patterns within the striatum, occupying specific subregions; for example, *Anxa1+* neurons preferentially project to the dorsal striatum, a region critical for motor control^6^. This supports the conclusion that these subtypes display divergent behaviors in motor control and reward processing^5^. However, our results also reveal that, despite their distinct projection patterns, these subtypes receive largely similar inputs across the brain, with some exceptions in cortical regions. This aligns with the idea that many dopamine neurons within a given region exhibit similar activity patterns, and suggests that their differential vulnerability is unlikely to be driven by differences in synaptic input. Still, several possibilities remain. One is that even subtle in input differences (for example, cortical regions in this case) may still have functional consequences. Alternatively, spatially distinct DAN subpopulations may receive inputs from different neuron groups within the same area, leading to distinct functions. Finally, SNc neurons may share broadly similar computational properties but display subtle differences in their response profiles. Their functional divergence may ultimately arise from how their projections interface with specific downstream cell types - for example, through the receptors expressed by their targets. A further layer of complexity emerges from the recurrent loop between the striatum and SNc: SNc dopaminergic neurons receive substantial inhibitory input from striatal medium spiny neurons while simultaneously projecting back to them^50,51^. The distinct projection profiles of dopaminergic neuron subtypes may therefore shape striatal dynamics in unique ways, with inhibitory feedback mechanisms subsequently fine-tuning dopaminergic signaling during diverse behavioral and learning processes. Additionally, a more detailed classification of the inputs and projections of each DAN subtype would be necessary to define specific input–output cell types, rather than the broader regional categories used here, as has been demonstrated in studies of fruit flies at the single-cell level^52^. For example, one of the major sources of input, the midbrain reticular nucleus (MRN), contains a heterogeneous mix of cell types, including glutamatergic and GABAergic neurons^53^. Unfortunately, our current methods do not allow us to determine which specific cell types contribute to the observed inputs. This granular approach may reveal intricate neurobiological dynamics obscured when broader classifications are used.

Finally, a recent study showed that *Anxa1+* neurons constitute a genetically distinct subset within the *Aldh1a1+* population and are functionally different from other dopaminergic subtypes, as demonstrated using their own transgenic line^5^. In parallel, we generated our own transgenic mouse line to manipulate these neurons with similar specificity. Although targeted manipulation of *Anxa1+* neurons led to significant bradykinesia and tremor similar to those observed in the early stages of disease in MitoPark mice, it did not affect immobile time or akinesia. These discrepancies might be explained by a dosage effect (i.e. akinesia requires the loss of more dopamine neurons than bradykinesia), or most likely by the involvement of other dopaminergic neuron subtypes that contribute to phenotypes emerging at later stages. For example, degeneration of *AGTR1+* or *Calb1+* neurons may also contribute to motor symptoms in the PD phenotype^8,54^. However, the motor deficits differ from those reported during *Anxa1+* neuron inhibition^48^. Thus, our study does not argue that *Anxa1+* neurons are the sole contributors to motor impairment. Rather, we show that their early loss is sufficient to generate motor deficits resembling those seen in early PD. In addition to the motor symptoms, we also found that inhibition of *Anxa1+* neurons produced a learning deficit. Given that cognitive dysfunction has been reported to precede motor symptoms in MitoPark mice^41^, these results suggest that *Anxa1+* neurons may also contribute to early non-motor symptoms as well.

However, given that other dopaminergic subtypes are also affected such as *Arhgap28+* neurons, our data do not exclude the possibility that additional cell types within the same group may play a role in learning. Further investigation will be important, particularly in relation to other early PD-associated features such as sleep behavior disturbances or constipation. The possibility of homeostatic compensation within DAN neural networks, with different neurons stepping in to sustain the functionality of behaviors affected by the loss of others, should also be discussed. We observed considerable overlap in the brain regions providing afferent connectivity to each dopamine subtype and this observation could be critical to understand the adaptability of these circuits. A critical approach to further unravel these complexities of adaptation as disease develops would involve recording whole-brain activity in PD model mice. Gaining a deeper understanding of the resilience and adaptability of neural circuits surrounding dopaminergic neurons will enhance our grasp of Parkinson’s disease and potentially foster innovative therapeutic strategies in the future.

## Materials and Methods

### Contact for reagents and resource sharing

All reagents and materials used in this study are listed in the Key Resources Table (DOI: 10.5281/zenodo.18273548). For additional information or requests, please contact the Lead Contact, Rui M. Costa.

### Mouse breeding and husbandry

All experimental protocols were performed according to National Institutes of Health (NIH) guidelines and in compliance with the regulations of the Institutional Animal Care and Use Committee at Columbia University. All experimental animals were 2 to 6 month-old mice housed on a 12 hr light/dark cycle with unrestricted access to food and water. Mice used for behavioral experiments were individually housed. The strain and lines used were: C57BL6/J (Jackson Laboratories, 000664), MitoPark mouse^13^, DAT-Cre mouse (Jackson Laboratories, 006660), VGlut2-Cre mouse (Jackson Laboratories, 028863), and Calb1-IRES2-Cre mouse (Jackson Laboratories, 028532). Anxa1-Cre (Jackson Laboratories, 040185) and DAT-Flp (Jackson Laboratories, 040184) mouse lines were developed and utilized in our lab (details are provided below)^35^. MitoPark mice (*DAT-Cre^+/-^;Tfam^loxP/loxP^*) were obtained by crossing *DAT-Cre^+/-^; Tfam^loxP/+^* mice with *Tfam^loxP/loxP^*mice to selectively knockout *Tfam* in dopaminergic neurons.

### Generation of transgenic mice (Anxa1-Cre and DAT-Flp)

Anxa1-Cre and DAT-Flp knock-in mice were generated by Cyagen (Santa Clara, CA) using CRISPR/Cas9. To generate Anxa1-Cre mice, one cell mouse embryos were microinjected with Cas9 protein, a gRNA (5’ GACATCCCAACTATTCTGCA-AGG 3’) targeting exon 13 in the vicinity of the stop codon, and a repair template comprised of homology arms and a ‘P2A-Cre-rBG pA’ cassette. Homology arms were generated by PCR from BAC clones RP23-53N8 and RP23-149B23 as template. Similarly, to generate DAT-Flp mice, a gRNA (5’ CCAACAGCCAATGGCGCAGC-TGG 3’) targeting exon 15, and a template with a ‘2A-FlpO-rBG pA’ cassette were used^35^. Homology arms were PCR amplified from BAC clones RP23-150M11 and RP23-34F24. Two silent mutations, aa 613L (CTG to TTA), and aa 614R (CGC to AGG) were introduced to prevent re-cleavage by Cas9 after homology-directed repair.

### Nuclear suspension, FACS and Single-nucleus RNAseq

We used procedures as previously described^55,56^. Briefly, MitoPark mice and their littermate controls (P56∼62 for 8 weeks, P112∼118 for 16 weeks and P168∼174 for 24 weeks; 4 female and 4 male at each time point) were anaesthetized with isoflurane and perfused with cold artificial cerebrospinal fluid (ACSF) containing N-methyl-d-glucamine (NMDG; 92 mM), KCl (2.5 mM), NaH2PO4 (1.2 mM), NaHCO3 (30 mM), HEPES (20 mM), D-Glucose (25 mM), Sodium L-ascorbate (5 mM), Sodium pyruvate (3 mM), MgSO4 (10mM), CaCl2 (0.5 mM), bubbled with carbogen gas (95% O2 and 5% CO2) at a pH of 7.2–7.4. The brain was sectioned at 400um using a vibratome (VT1200S, Leica Microsystems) on ice, and the ventral midbrain (SNc and VTA) were microdissected from three consecutive sections (−2.48 to −3.88mm from Bregma; Franklin and Paxinos, 2008). The tissues were transferred to microcentrifuge tubes, flash frozen in dry ice with ethanol, and stored at −80C until later use. For nuclei isolation^22^, frozen tissues were placed into a homogenization buffer that consisted of 10mM Tris pH 8.0, 250mM Sucrose, 25mM KCl, 5mM MgCl2, 0.1% Triton-X 100, 1X Protease inhibitor (Promega G6521), 0.5% RNasin Plus RNase inhibitor (Promega N2615) and 0.1mM DTT (Promega P1171). Tissues were placed into a 2ml dounce homogenizer (Sigma D8938) and homogenized using 10 strokes of the loose dounce pestle followed by 10 strokes of the tight pestle to liberate nuclei. Homogenate was strained through a 30μm cell strainer (Miltenyi Biotech 130-098-458) and centrifuged at 900xg for 10 minutes to pellet nuclei. Nuclei were then resuspended in blocking buffer containing 1X PBS supplemented with 0.8% nuclease-free BSA (Sigma, OmniPur 2905) and 0.5% RNasin Plus RNase inhibitor. Prior to fluorescence-activated nuclei sorting (FACS), DAPI (Sigma, D9542) was applied to nuclei suspensions at a final concentration of 0.1μg/ml and nuclei suspensions were filtered through a 30μm nylon mesh (Sysmex, 04-004-2326) to remove aggregates. FACS was performed at the Zuckerman Institute Flow Cytometry platform using a MoFlo Astrios EQ (Beckman Coulter) sorter. Single nuclei were captured by gating DAPI+ events, excluding debris and doublets. Both cell suspension and collection chamber were kept at 4C during sorting. Sorted nuclei were spun down immediately and barcoded using 10x Genomics Chromium Single Cell 3’ Reagent Kit v3 according to the manufacturer’s protocol. Samples were processed and libraries were prepared/sequenced by the Columbia JP Sulzberger Genome Center Single Cell Analysis Core.

### RNAseq data processing and analysis

Sequencing data were aligned and quantified using the Cell Ranger Single-Cell Software Suite (v.6.1.2) with default parameters against the ENSEMBL GRCm38/mm10 reference genome^57^. The pre-filtered datasets were then imported into the R Seurat package for downstream analysis (v5.0.2)^58^. We first removed low-quality cells (cell containing < 200 genes and genes expressed in <3 cells of the data). After the filtering, contaminating ambient RNA reads were filtered by using SoupX^59^. Doublets were predicted using the DoubletFinder R package^60^, and were excluded for downstream analysis. After quality control and cell filtering (1000<nFeature_RNA<7500; nCount_RNA<40000; percent_mt<1, percent_ribo<1, percent_hemo<1), the datasets were processed for identifying cell populations by Seurat pipeline. Briefly, pre-processed gene expression matrices were normalized by the NormalizedData function. A total of 2,000 most variable features were then selected with the FindVariableFeatures and scaled by ScaleData command (while regressing out mitochondrial genes) for the Principal Component Analysis (PCA), which was performed by the RunPCA command for 30 PCs. After integration with Harmony^61^, the data was subsequently processed with Louvain clustering and dimensionality reduction via Uniform Manifold Approximation and Projection (UMAP) using 30 principal components, reduction = “harmony”, and resolution of 0.8, to visualize nuclear transcriptomic profiles in two-dimensional space. For annotation, we used the Allen Brain Cell (ABC) Atlas tool MapMyCells (RRID:SCR_024672, the reference taxonomy: 10x Whole Mouse Brain (CCN20230722), mapping algorithm: hierarchical mapping)^23^ to assign neuronal classes or subclasses to nuclei from a whole dataset. To validate the classification, we examined the clusters using well-established markers such as *Th*, *Slc17a6* (Vglut2), and *Slc32a1* (Vgat), which confirmed that the annotation was reasonable. To subset the dopaminergic population, we first used the annotation (21_MB_Dopa) to identify and extract the relevant population. We then re-scaled the extracted data and applied the supertype or cluster annotation to the subset and performed further analysis. Differentially expressed genes in each cluster were identified by Wilcoxon test (the FindAllMarkers function in Seurat),

### Differential cell abundance

To identify differentially abundant cell populations in each dataset, we used two methods: MASC (version 0.10)^24^ and Milo (version 1.99)^30^. MASC is a single cell association method for testing whether a certain status influences the membership of single cells while accounting for technical confounds and biological variation. For all datasets, we included sex as a fixed effect, and individual as a random effect in the model. We then tested the significance of association between the status (i.e. MitoPark mice and their littermate controls at each stage, or MitoPark mice at different stages) with the clusters. Cell subpopulations were considered significantly associated with a status at FDR-adjusted P<0.05 and absolute odds ratio >0. Milo constructs local neighborhoods (k=30, d=30) of transcriptionally similar cells on a k-nearest neighbor (kNN) graph derived from the single-cell dataset, effectively grouping small clusters of similar cells together. Each neighborhood represents a subset of cells with comparable transcriptomic profiles and may correspond to fine-grained cellular subpopulations. Milo then quantifies the number of cells from each sample within each neighborhood and compares their relative abundances across experimental conditions, for example, assessing whether certain neighborhoods are enriched in diseased samples relative to healthy controls. Statistical testing is performed to determine whether these differences are significant or attributable to random variation. By systematically applying this procedure across all neighborhoods, Milo identifies cellular subpopulations that exhibit condition-specific changes in abundance.

### Gene ontology enrichment analysis

We used the g:Profiler web tool and gprofiler2 R package (version 0.2.3)^62^ to identify significantly enriched gene ontology (GO) terms from upregulated or downregulated gene lists in dopaminergic clusters.

### Stereotaxic viral injections

Before starting the surgery mice were subcutaneously injected with Buprenorphine XR (0.5–1 mg per kg body weight). The mouse head was shaved, cleaned with 70% alcohol and iodine, an intradermic injection of bupivacaine (2 mg per kg body weight) was administered, and a small incision from anterior to posterior was made on the skin to allow for aligning the head and drilling the hole for the injection site. Surgeries were performed under sterile conditions and isoflurane (1%–5%, plus oxygen at 1-1.5 l/min) anesthesia on a motorized stereotactic frame (David Kopf Instruments, Model 900SD). Throughout each surgery, mouse body temperature was maintained at 37C using an animal temperature controller (ATC2000, World Precision Instruments). For anterograde tracing experiments, animal were unilaterally injected with 300nL of AAV5-CAG-FLEX-GFP (titer: 4.5E12 vg/mL; UNC Vector Core) into the right hemisphere of the substantia nigra pars compacta (SNc; AP −3.16 mm, ML 1.3 mm, DV −4.0 mm) using a Nanoject III Injector (Drummond Scientific, USA) at a pulse rate of 5.1 nL per second, 20 nL per pulse every 5 s. After injection, the pipette was held in place for 5 minutes before raising to ensure minimize viral efflux. For whole-brain retrograde tracing experiments using rabies virus, 100nL of a 2:1 mixture of the helper viruses, AAV1-CAG-FLEX-N2cG-mKate2.0 (titer: 9.83E12 vg/mL; Janelia Viral Core) or AAVDJ-CAG-FlpX-N2cG-mKate2.0 (titer: 2.60E12 vg/mL; Zuckerman Institute Vector Core) and AAV5-Esyn-DIO-TVA950-EYFP-WPRE (titer: 1.44E13 vg/mL; Salk Viral Vector Core), was injected into the SNc. Two weeks later, 300nl of EnvA-N2c-deltaG-tdTomato (titer: 1.5E9 ffu/ml; Center for Neuroanatomy with Neurotropic Viruses, CNNV) was injected within the same area at a 10-degree angle (SNc; AP −3.16 mm, ML 2.0mm, DV −4.0 mm) to prevent labeling the injection tract with TVA. For monosynaptic labeling from the primary motor cortex or the dorsal striatum, 100nL of a 2:1 mixture of the helper viruses, AAV1-CAG-FLEX-N2cG-mKate2.0 (titer: 9.83E12 vg/mL; Janelia Viral Core) and AAV5-Esyn-DIO-TVA950-EYFP-WPRE (titer: 1.44E13 vg/mL; Salk Viral Vector Core), was injected into the SNc. 100nL of AAVDJ-Ef1a-fDIO-mScarlet (titer: 6.30E12 vg/mL; Zuckerman Institute Vector Core) was injected in the motor cortex or the striatum. Two weeks later, 300nl of EnvA-N2c-deltaG-H2B-HA-FlpO (titer: 1.0E9 ffu/ml; Zuckerman Institute Vector Core) was injected within the same area at a 10-degree angle (SNc; AP −3.16 mm, ML 2.0mm, DV −4.0 mm). After the injection, the skull was cleaned and the skin sealed with sutures and Vetbond tissue adhesive (3M, Maplewood, MN, USA). For ablation experiments, 150 nL of AAV5-EF1a-FLEX-taCasp3-TEVp virus (titer 4.20E12 vg/mL; UNC Vector Core) was bilaterally injected into the SNc of DAT-Cre at two specific coordinates (SNc-medial: AP −3.16 mm, ML 1.10 mm, DV −4.1 mm; SNc-lateral: AP −3.16 mm, ML 1.50 mm, DV −4.1 mm). These coordinates effectively targeted and ablated SNc dopaminergic neurons. For silencing neurotransmission, 300nL of AAVDJ-hSyn-FLEX-TeLC-EYFP (titer 7.40E12 vg/mL; Zuckerman Institute Vector Core) was bilaterally injected in the SNc of Anxa1-Cre (SNc; AP −3.16 mm, ML 1.3 mm, DV −4.0 mm).

### Open field and cylinder test behavior assays

Mice were habituated with a wireless motion sensor in a square arena (40 x 40 x 20 cm, length x width x height) housed inside a sound-attenuating chamber for 30 minutes on the first day. Imaging was conducted using a 1280 x 1024 pixel FLIR camera (Point Grey Flea) with a CS mount lens, recording at 30 Hz. For the open-field test, mice were imaged for 30 minutes from a top view, and for the cylinder test (a cylinder arena: 15cm diameter x 30cm length), they were imaged for 15 minutes from a side view. Video acquisition was performed using the commercial software FlyCapture and the open-source visual language Bonsai^63^. Mouse centroid coordinates (i.e., spine2 in DeepLabCut tracking, details provided below) during the open-field recording were used to calculate basic measures of activity (e.g., trajectory or velocity). Acceleration data was collected at a sampling rate of 200 Hz (details provided below).

### Pose estimation and tracking analysis

DeepLabCut^64^ was used to estimate the pose of the mice and track their movements throughout the trials. In the open-field test, seven body parts (nose, right ear, left ear, spine1, spine2, spine3, tail) were labeled to create a skeleton for pose estimation. For the cylinder test, two body parts (motion sensor and nose) were labeled. A total of 1,100 frames were labeled for the open-field test (training: 1.03 million iterations, training error: 1.72 pixels, test error: 2.76 pixels), and 1,040 frames were labeled for the cylinder test (training: 1.03 million iterations, training error: 2.31 pixels, test error: 6.8 pixels). The pose data was filtered using a median filter and verified through visual inspection. For the analysis in the open-field test, to ensure consistent data, we only used data from the 5 to 20-minute mark (15 minutes total) within a 30-minute recording, as mice tend to move differently during the first few minutes after being placed from their home cage into the field. Immobility was defined as any tracking point with a speed of less than 0.5 cm/s for a duration of more than 0.5 seconds, and estimates with a likelihood lower than 0.95 were excluded. Average speed was calculated only during periods when the animal was mobile. In the cylinder test, the full 15 minutes of data were used for analysis, since mice often exhibit exploratory rearing behaviors during the first few minutes. For quantifying the number of rears, we tracked the vertical positions of the nose and motion sensor on the mouse head using DeepLabCut. If the tracking was sustained for at least 0.5 seconds in the same position, it was defined as a single rearing behavior. The height thresholds were determined using DeepLabCut’s output (i.e., filtered trajectory plots), and the number of rears instances was recorded.

### Motion sensor acceleration data analysis

Wireless motion sensor data was collected using the WEAR system developed by the Champalimaud Hardware Platform. The sensor is compact and lightweight (∼1.8 g), capable of sampling nine-axis motion data from a three-axis accelerometer, gyroscope, and magnetometer at rates up to 200 Hz^15^. This device communicates with computers through a base station designed by the platform, utilizing the Harp system. The base station is accessible via a software GUI (e.g., Harp Wear), allowing for easy adjustment of sensor parameters to best fit experimental needs. Additionally, it is compatible with Bonsai software^63^, enabling synchronization of the WEAR sensor data with other data sources, such as cameras. The data was first filtered using a 5^th^ order, 1Hz low-pass Butterworth filter to separate the gravitational acceleration component of each axis. This component was then subtracted to the original acceleration data to obtain the fast acceleration changes induced by mice’s movements (body acceleration). Total body acceleration was determined by calculating the vector norm of the three body acceleration axis. To assess the emergence of rhythmic, oscillatory movements, we performed a spectral analysis of the body’s dorso-ventral acceleration axis. We used the signal.spectrogram function of the Scipy python library^65^ to calculate a spectrogram with consecutive Fourier transforms, using segments of 256 samples with a 50% overlap. The spectrogram was normalized by transforming the power values of each window to a 0-1 range (maximum= 1 and minimum=0). For each mouse a final measure was obtained by calculating the median normalized power considering all the windows of the spectrogram.

### Pavlovian conditioning

Mice were habituated to custom-made Pavlovian chambers (17 x 14 x 20 cm, length x width x height) placed inside sound-attenuating boxes. They were food-restricted throughout the training period and maintained at 85% of their original body weight. Each chamber was equipped with a nose poke port featuring an infrared beam and stimulus LED, as well as a speaker. A food reward (6-8 µl of 10% sucrose solution) was delivered through a magazine spout equipped with a lickometer. Cues and reward delivery were controlled by a Python-based framework (i.e., the pyControl system^66^), which also recorded all behavioral timestamps. Behavior was monitored using top (USB camera, ELP, China) and side (Flea3, FLIR, USA) cameras, with video capture managed by FlyCapture and Bonsai softwares. For stimuli, a 10-second LED light served as the CS+, while a 10-second tone (4 kHz, 75 dB) served as the CS-. At the onset of each CS+, a drop of sucrose solution was delivered to the tip of the spout, whereas the CS- was not paired with a reward. CS+ and CS- were presented independently on a random inter-trial interval schedule of 120 seconds (ITI = RI-120). A training session concluded once the animal received a total of 30 rewards. Each animal underwent one training session per day for five consecutive days. Learning progress was assessed by monitoring the lick rate and the latency for each mouse to make its first lick after the onset of each CS+.

### Histology

Mice were deeply anesthetized with isoflurane and transcardially perfused with PBS, followed by ice-cold 4% paraformaldehyde. Brains were then removed for histological analysis. Coronal sections were cut at 30 µm for immunostaining or 75 µm for 3D reconstruction using a Leica VT1000 vibratome. The tissue was rinsed twice in 1x PBS and then permeabilized in PBS containing 0.4% Triton X-100 (PBST). For quantification of *Th+* cells in MitoPark mice and Anxa1-Cre mouse line, immunohistochemistry was performed with primary antibodies by incubating the sections with an anti-GFP antibody (Chicken, Aves labs Inc, GFP-1020) and a TH antibody (Mouse, ImmunoStar, 22941) diluted at 1:2000 in 0.4% Triton X-100 PBS (PBST) overnight at 4C. To visualize the dorsal-tier or ventral-tier of the SNc DANs, we also used a Calbindin D28K (Rabbit, Synaptic Systems, 214002) or an Aldh1a1 antibody (Goat, R&D Systems, AF5869). The sections were then incubated with secondary antibodies (Donkey anti-Chicken Alexa Fluor 488, JacksonImmunoResearch, 703-545-155; Donkey anti-Goat Alexa Fluor 568, Invitrogen, A11057; Donkey anti-Rabbit Alexa Fluor 568 (Invitrogen, A10042); Donkey anti-Mouse Alexa Fluor 647, Invitrogen, A31571) diluted at 1:2000 in 0.4% PBST overnight at 4C. DAPI (Sigma D9542; 1:1000) was used as a counterstain in all experiments.

### Image acquisition, processing, and analysis

Coronal sections (30µm or 75μm) were serially mounted on slides and sections were imaged using an automated slide scanner (Nikon AZ100 Multizoom microscope) equipped with a 4x 0.4NA Plan Apo objective (Nikon Instruments Inc) and P200 slide loader (Prior Scientific), controlled by NIS-Elements using custom acquisition scripts (Nikon Instruments Inc.). Image processing and analysis using BrainJ proceeded as previously described^21^. Briefly, brain sections were aligned and registered using two-dimensional (2D) rigid-body registration. After background subtraction ilastik was used to detect cell bodies and neuronal processes^67^. Probability images for class (soma and neuronal processes) were generated using the pixel classification approach in ilastik, ensuring improved accuracy for segmentation over fluorescence intensity alone. To map the location of these structures to an annotated brain atlas, serial tissue sections were registered into a single volume before 3D registration to the template brain (Allen Brain Atlas Common Coordinate Framework) using Elastix^68^. Initial measurements and data visualization were performed in ImageJ, and subsequent analyses were performed in Python.

### Statistics and reproducibility

All statistical analyses were conducted using R or Python. Data are presented as mean with 95% confidence intervals (CI) or as the standard error of the mean (SEM), unless otherwise specified. Statistical significance was assessed using a two-sided Mann–Whitney U test, and for multiple comparisons, a two-way ANOVA followed by Tukey’s HSD test was applied. To further validate the results, a linear mixed-effects model (LMM) with Tukey’s post hoc comparisons was also conducted for certain results (*p < 0.05, **p < 0.01, ***p < 0.001, ****p < 0.0001). All measurements were obtained from distinct, independent samples unless otherwise stated.

### Data and code availability

All data, code, experimental protocols, and essential laboratory materials used or generated in this study are listed in the Key Resources Table, together with their corresponding persistent identifiers at DOI: 10.5281/zenodo.18273548.

## Acknowledgments

We would like to thank Zuckerman Institute’s Cellular Imaging platform for providing instrument use and technical advice. Special thanks to Gabriela J. Martins, Mariana L. Correia and Helena M Seuffert for mouse maintenance and Drew Baughman, Mafalda Vicente and Chrissy Weber-Schmidt for additional support with lab and mouse management. We also appreciate Helio Rodrigues for designing and constructing behavioral equipment, and Kim Ritola for creating custom viral constructs. Our gratitude extends to Ira Schieren for assisting with FACS sorting. This work was supported by grants from the Aligning Science Across Parkinson’s (ASAP-020551) through the Michael J. Fox Foundation for Parkinson’s Research (MJFF), the Parkinson’s Disease Foundation (PDFPF-RCE-1948), the National Institutes of Health (NIH, U19: 5U19NS104649-03), and a NARSAD Young Investigator Grant from the Brain & Behavior Research Foundation (30086). This research was funded in part through the NIH/NCI Cancer Center Support Grant P30CA013696 and used the Genomics and High Throughput Screening Shared Resource. Additional support was received from the National Center for Advancing Translational Sciences, NIH, under Grant Number UL1TR001873. Rabies viruses were produced by the Center for Neuroanatomy with Neurotropic Viruses (CNNV), supported by the NIH under Grant P40 OD010996. For the purpose of open access, the author has applied a CC BY public copyright license to all Author Accepted Manuscripts arising from this submission.

## Author contributions

A.F. and R.M.C. designed the study, interpreted the results, and wrote the manuscript. A.F. conducted the experiments and analyzed the data. D.N. generated viral constructs and assisted with the production of transgenic mice. Z.R.L., A.Y., B.T., and V.M. contributed to transcriptome data analysis and interpretation of sequencing results. L.A.H. developed the automated pipeline for anatomical reconstruction and analysis, and contributed to histological assessments. C.W. produced the rabies virus. T.S. and J.A.d.S. assisted with accelerometer experiments and data analysis. R.M.C. supervised the project. All authors contributed to the interpretation of results and manuscript editing.

---

**Extended Data Fig. 1.**
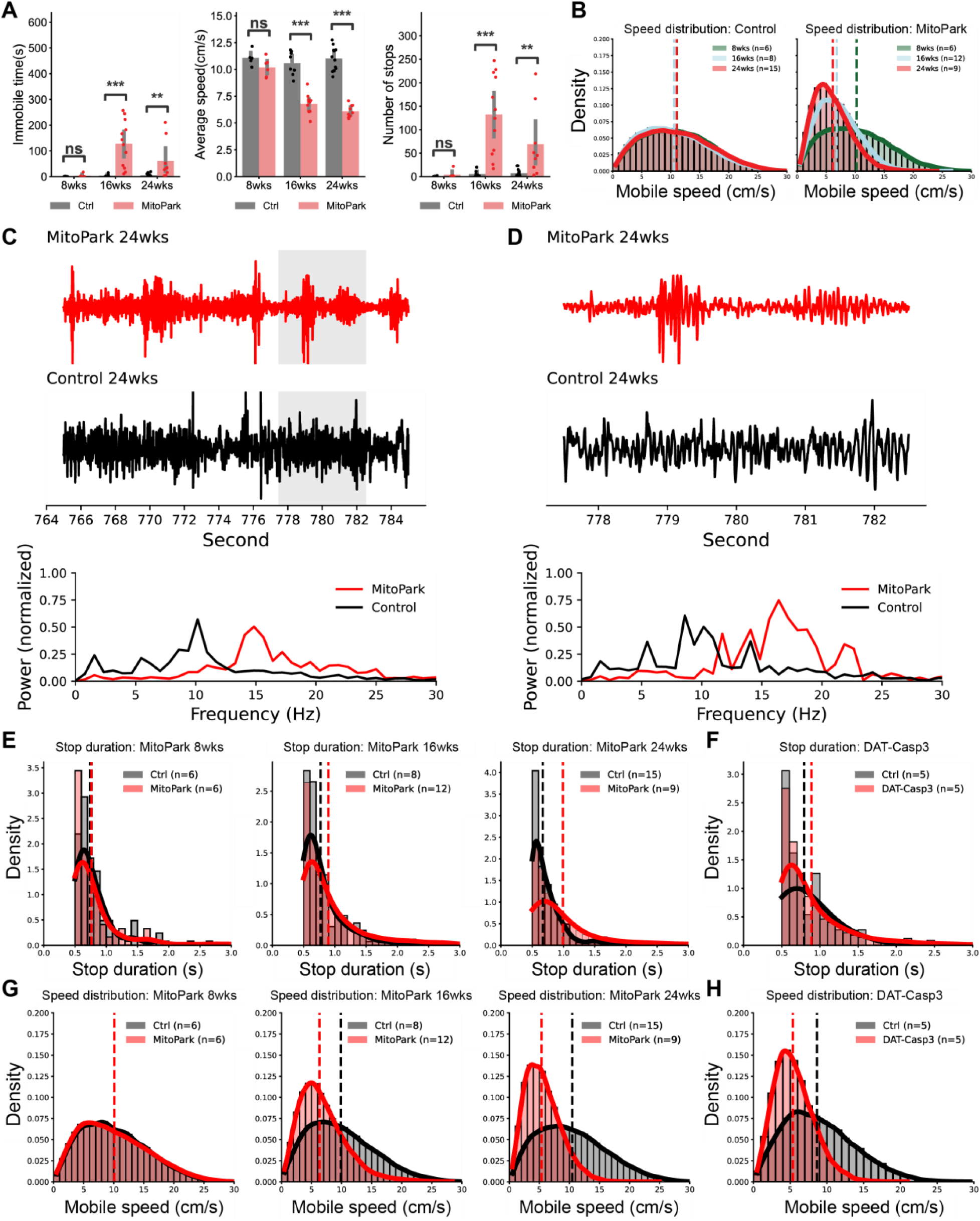
Behavioral changes associated with the progression of neurodegeneration in MitoPark mice, Related to Fig. 1. **A-B.** Environmental novelty does not influence basic locomotor activity in MitoPark animals. Measures of immobility time, average speed, and number of stops on the initial habituation day (Day 0) did not differ significantly from those recorded on the actual testing day (Day 1). Although average speed was slightly higher on Day 0, this likely reflects heightened exploratory drive and motivation in response to the novel environment (A). Similarly, speed distribution analysis showed no significant differences between the two days (B). **C-D.** Examples of raw traces of dynamic acceleration in MitoPark animal 24 weeks (red) and a control animal (black), with an enlarged view (gray square in A) shown in **C**. MitoPark mice often show specific 12-18Hz oscillations. **E-H.** These figures show the probability density functions (PDFs) for stop duration and speed, with duration (s) and speed (cm/s) on the x-axis and probability density on the y-axis. In MitoPark mice, stop durations increased and speed distributions shifted leftward at later stages (E, G). DAT-Casp3 animals exhibited a similar pattern, with prolonged stop durations and reduced speeds (F, H), resembling the later-stage phenotype of MitoPark mice.

**Extended Data Fig. 2.**
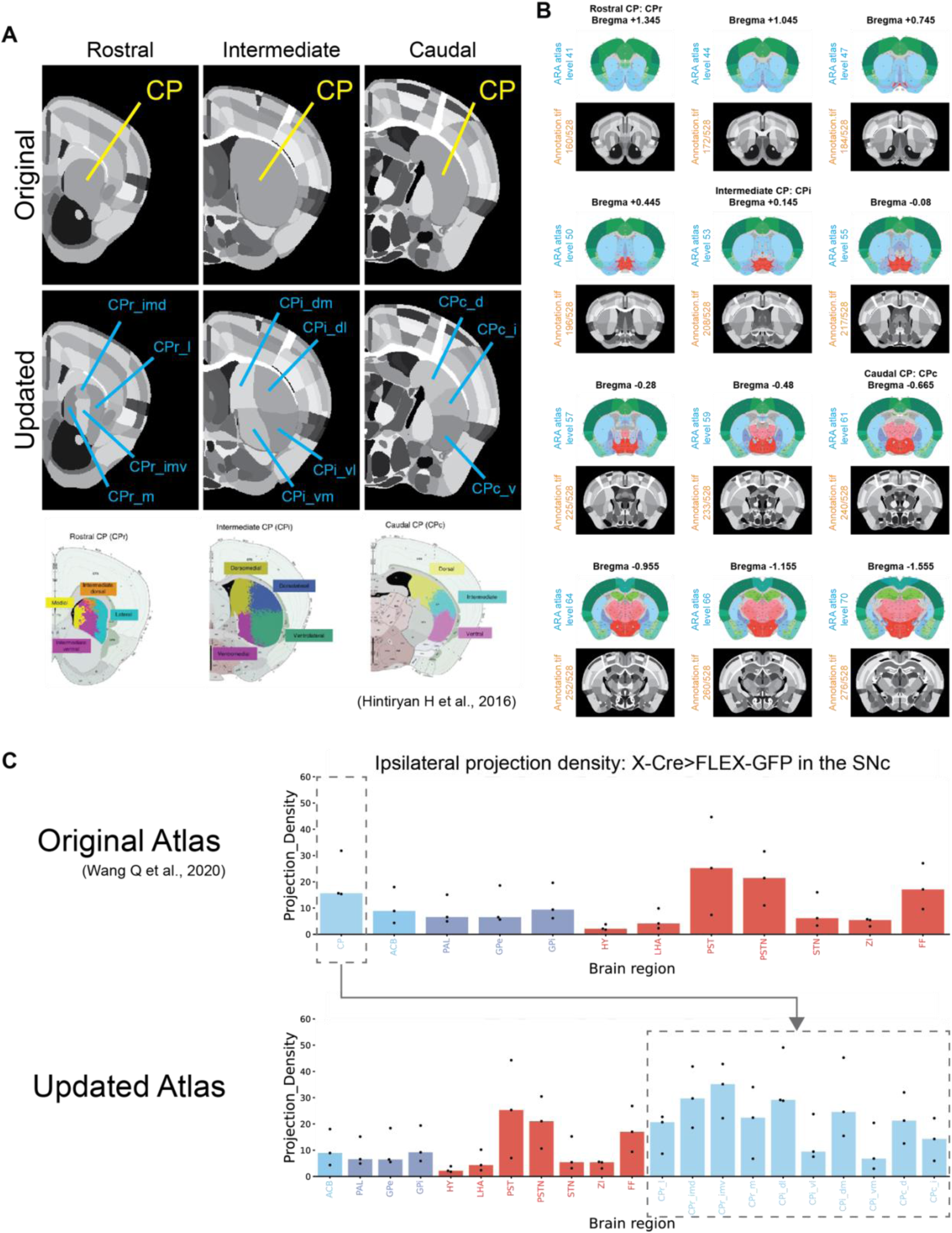
An updated atlas incorporates striatal subregions and enhances the analysis of cells and projections, Related to Fig. 2 and Fig. 5. **A.** Top: The original Allen Common Coordinate Framework (Allen CCF; version 3 released in 2017). Middle: An updated version of the Allen CCF, incorporating subregions of the striatum (CP: caudate putamen) based on the cortical projections shown at the bottom (CPr_imd: Rostral CP – Intermediate dorsal, CPr_l: Rostral CP – Lateral, CPr_imv: Rostral CP – Intermediate ventral, CPr_m: Rostral CP – Medial, CPi_dm: Intermediate CP – Dorsomedial, CPi_dl: Intermediate CP – Dorsolateral, CPi_vl: Intermediate CP – Ventrolateral, CPi_vm: Intermeidiate CP – Ventromedial, CPc_d: Caudal CP – Dorsal, CPc_i: Caudal CP – Intermediate, CPc_v: Caudal CP – Ventral). **B.** Alignment of coordinates between the ABA (Allen Brain Atlas) and Allen CCFv3, with manually segmented subregions of the striatum. **C.** Comparison of projection density of dopamine neurons in SNc using the original atlas versus the updated atlas. The original atlas (top) does not include subdivisions of the caudate-putamen (CP), while the updated atlas (bottom) defines 11 subregions based on cortical projections, providing a more detailed representation of local projections within the striatum.

**Extended Data Fig. 3.**
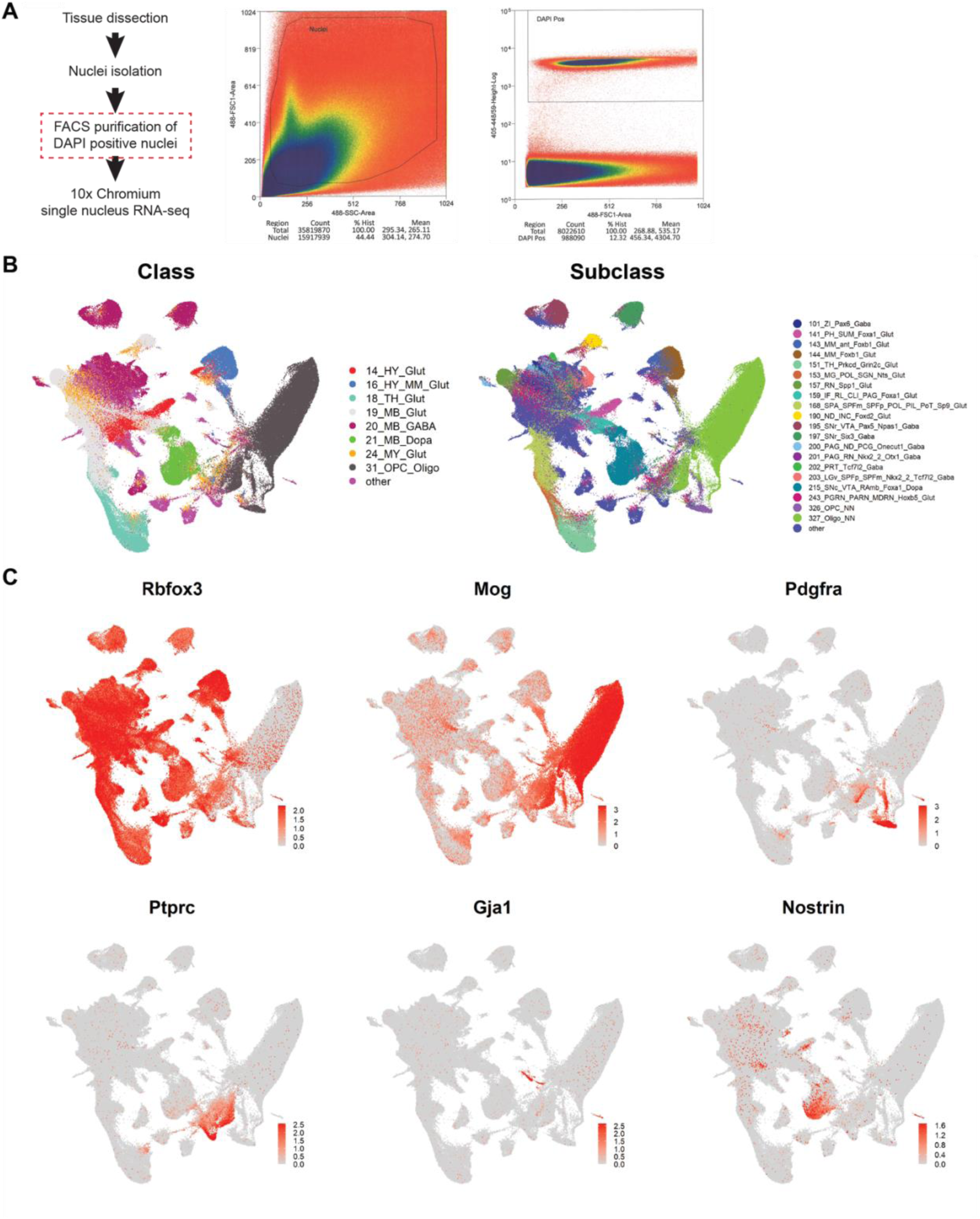
The integrated dataset with the Allen Brain Cell (ABC) Atlas, Related to Fig. 2. **A.** UMAP plots showing all nuclei in the ventral midbrain of MitoPark mice and their controls with the class (left) or subclass (right) annotation of the ABC Atlas. Dopamine cluster is classified as Class: 21_MB_Dopa or Subclass: 215_SNc_VTA_RAmb_Foxa1_Dopa. **B.** Representative feature plots to label main class of cells. Neuronal cells, *Rbfox3*; Oligodendrocytes, *Mog*; Oligodendrocyte precursor cells (OPCs), *Pdgfra*; Microglia, *Ptprc*; Astrocytes, *Gja*; Endothelial cells, *Nostrin*.

**Extended Data Fig. 4.**
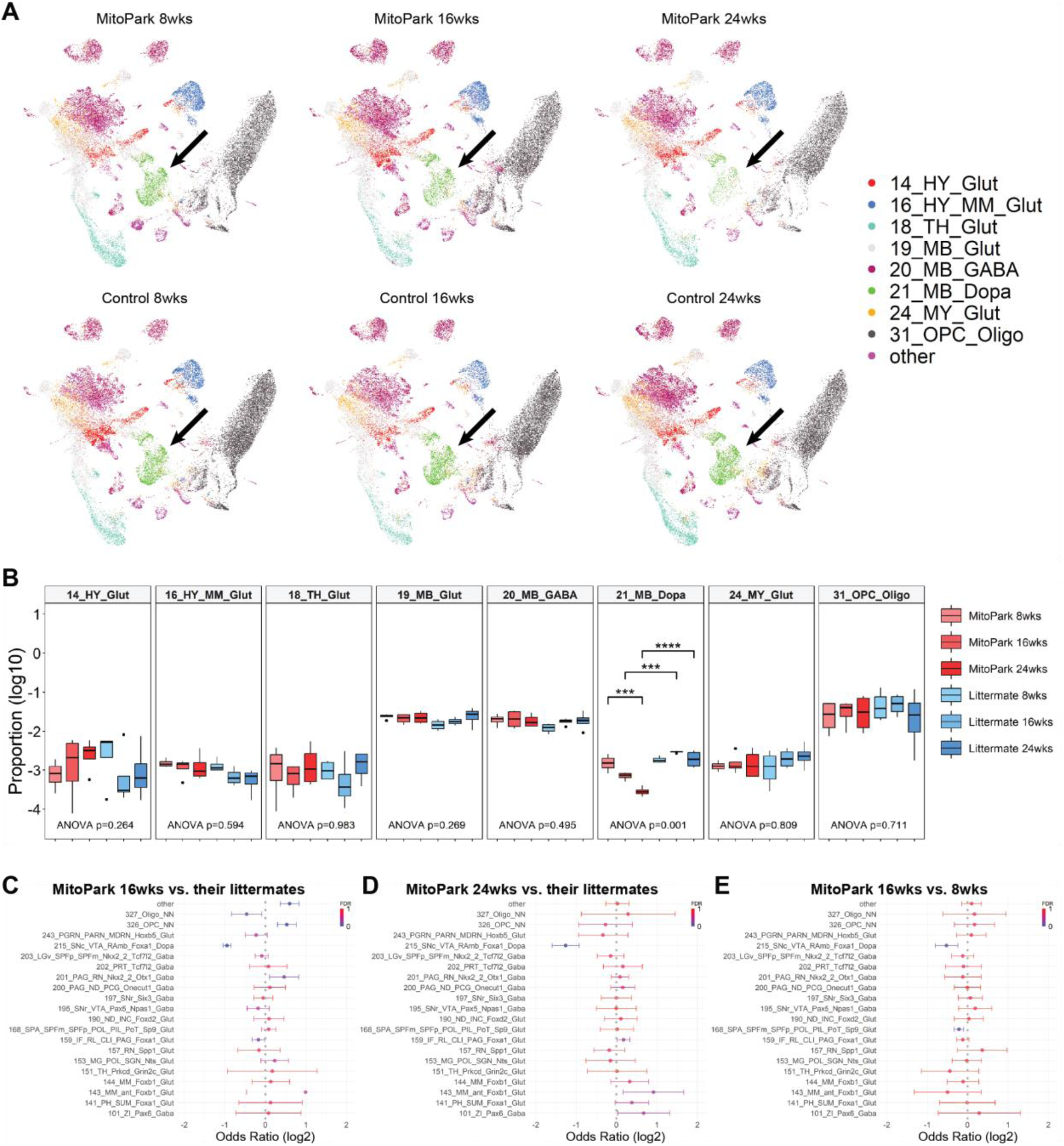
A significant reduction in DANs was observed in the later weeks of MitoPark mice, Related to Fig. 2. **A.** UMAP plots showing all nuclei in the ventral midbrain of MitoPark mice and their controls with the class annotation of the ABC Atlas. Dopamine cluster (black arrow) becomes visibly thinner than other cell types in MitoPark mice. **B.** Box and whisker plots showing the fraction of major cell types originating from the ventral midbrain of MitoPark mice, their littermate controls and B6 WT mice (each n=8), with plot center, box and whiskers corresponding to median, interquartile range (IQR) and 1.5xIQR, respectively. MitoPark mice in the later stages exhibited a significant reduction in the proportion of DANs (21_MB_Dopa), without affecting other cell types. Statistical significance was determined by two-way ANOVA (p-value indicates the interaction effect between genotype and age) followed by Tukey’s HSD test for multiple comparisons (*p < 0.05, **p < 0.01, ***p < 0.001, ****p < 0.0001). **C-E.** Odds-ratio estimated of major cell types annotated by subclass. **(C)** MitoPark 16 weeks versus their littermate controls; 215_SNc_VTA_RAmb_Foxa1_Dopa (OR=-0.95, FDR-adjusted P<0.05). **(D)** MitoPark 24 weeks versus their littermate controls; 215_SNc_VTA_RAmb_Foxa1_Dopa (OR=-1.26, FDR-adjusted P<0.05). **(E)** MitoPark 16 weeks versus 8 weeks; 215_SNc_VTA_RAmb_Foxa1_Dopa (OR=-0.51, FDR-adjusted P=0.086).

**Extended Data Fig. 5.**
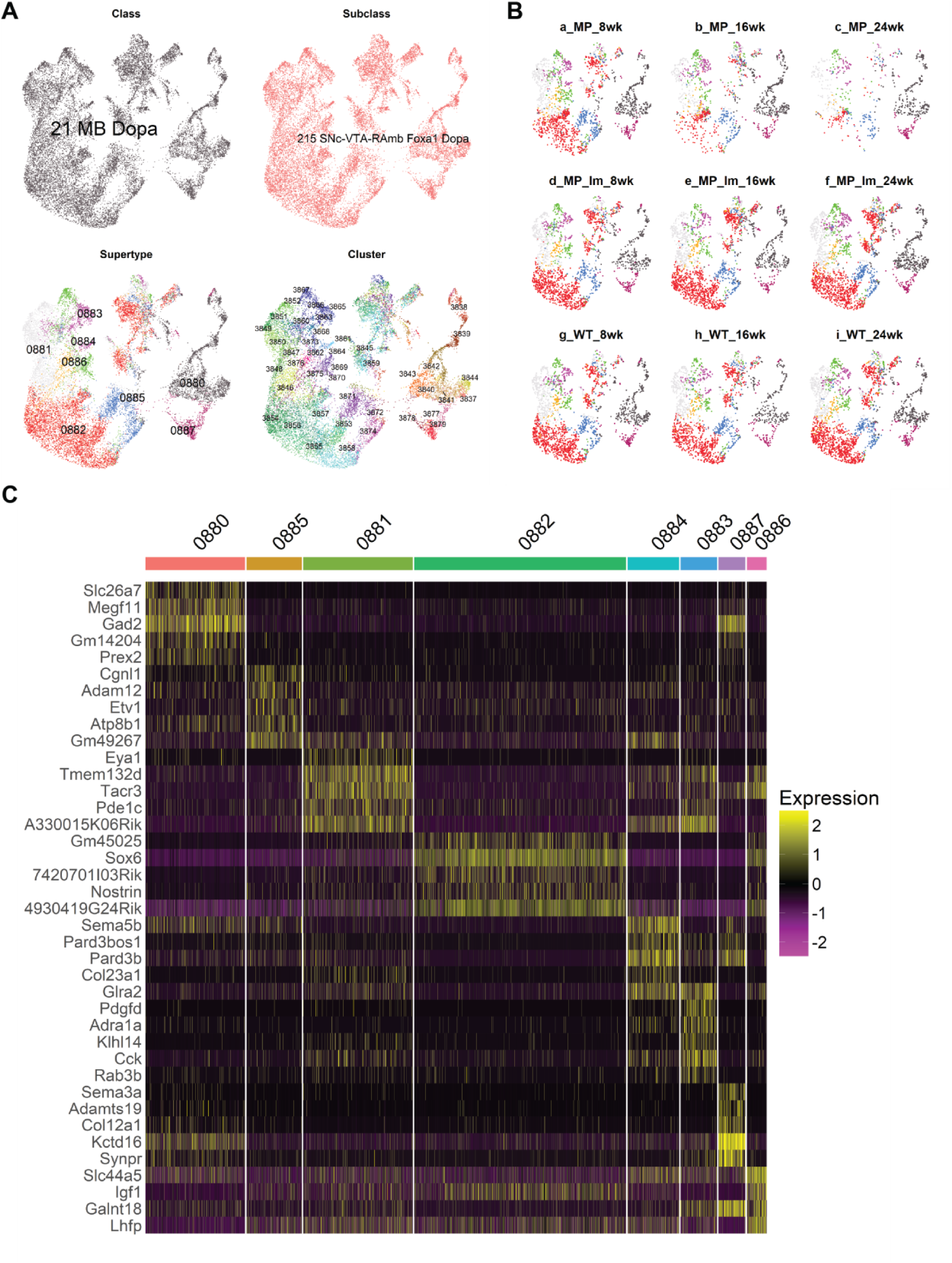
The dopaminergic cluster with distinct marker genes, Related to Fig. 3. **A.** UMAP plots showing dopamine nuclei in the MitoPark mice and their controls with the annotations (Class, Subclass, Supertype and Cluster) of the ABC Atlas. **B.** UMAP plots illustrating the distribution of dopamine nuclei, categorized by supertype annotation based on the species and age. **C.** Top five marker genes for each dopamine neuron subtype categorized by supertype annotation.

**Extended Data Fig. 6.**
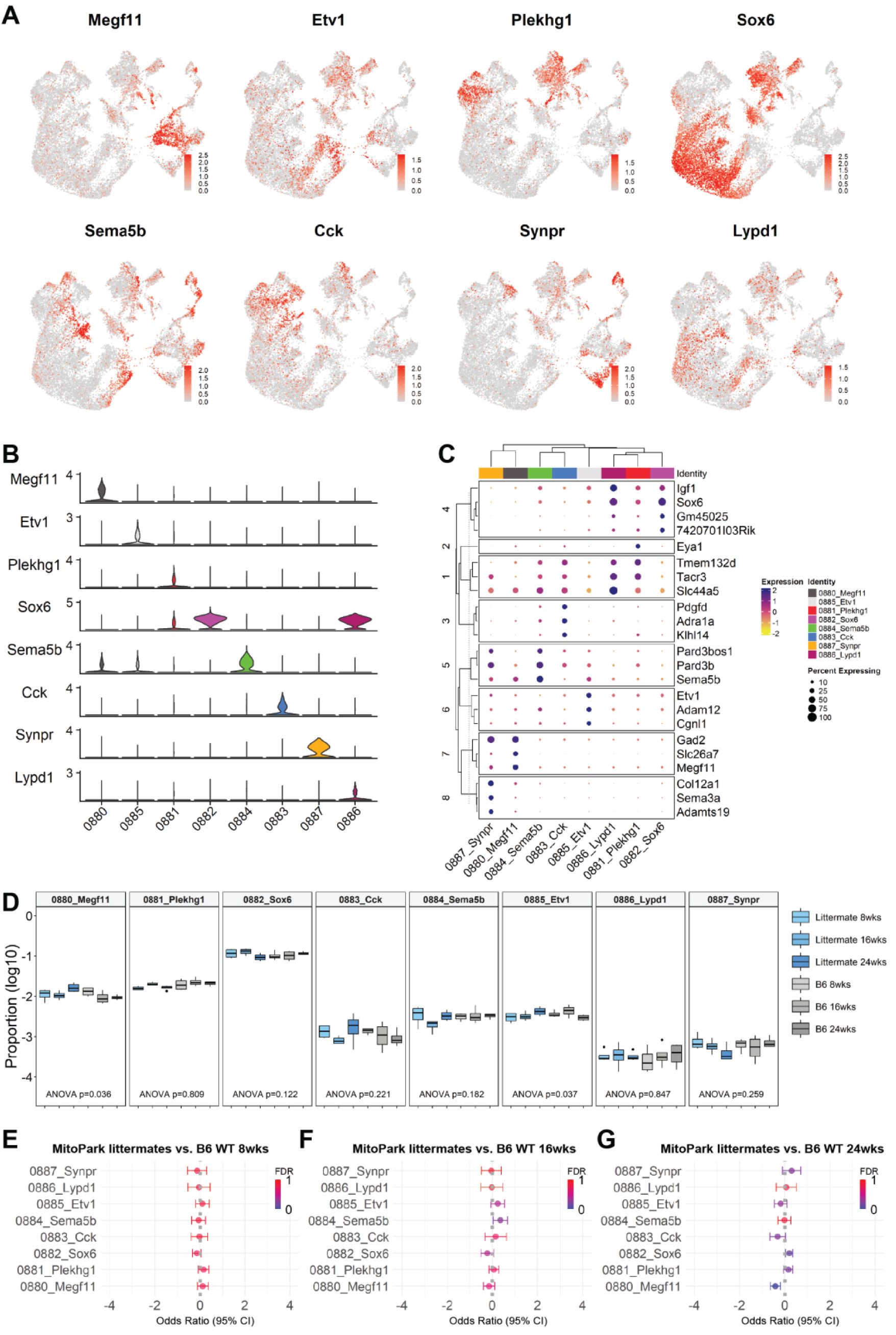
Marker feature expression in the dopaminergic cluster, Related to Fig. 3. **A.** Feature plots depicting the expression of marker genes across dopamine clusters, classified according to supertype annotations.**B.** Violin plot indicating the expression distribution of the top marker gene of each cluster. **C.** Clustered dot plot profiling of marker genes for each dopamine cluster. X-axis represents the dopamine clusters, and the Y-axis shows the top three marker genes. Dot diameter indicates the proportion of cluster nuclei expressing a given gene. The color scale represents the gene expression levels, with black indicating high expression and white indicating low expression. **D.** Distribution of dopamine subtype proportions across MitoPark littermate control animals and B6 WT animals. **E-G.** There is no significant age-related degeneration in dopamine populations. **(E)** MitoPark littermate controls versus B6 WT animals at 8 weeks; 0882_Sox6 (OR=-0.14, FDR-adjusted P=0.80). **(E)** MitoPark littermate controls versus B6 WT animals at 16 weeks; 0882_Sox6 (OR=-0.21, FDR-adjusted P<0.41). **(F)** MitoPark littermate controls versus B6 WT animals at 24 weeks; 0882_Sox6 (OR=0.20, FDR-adjusted P<0.11).

**Extended Data Fig. 7.**
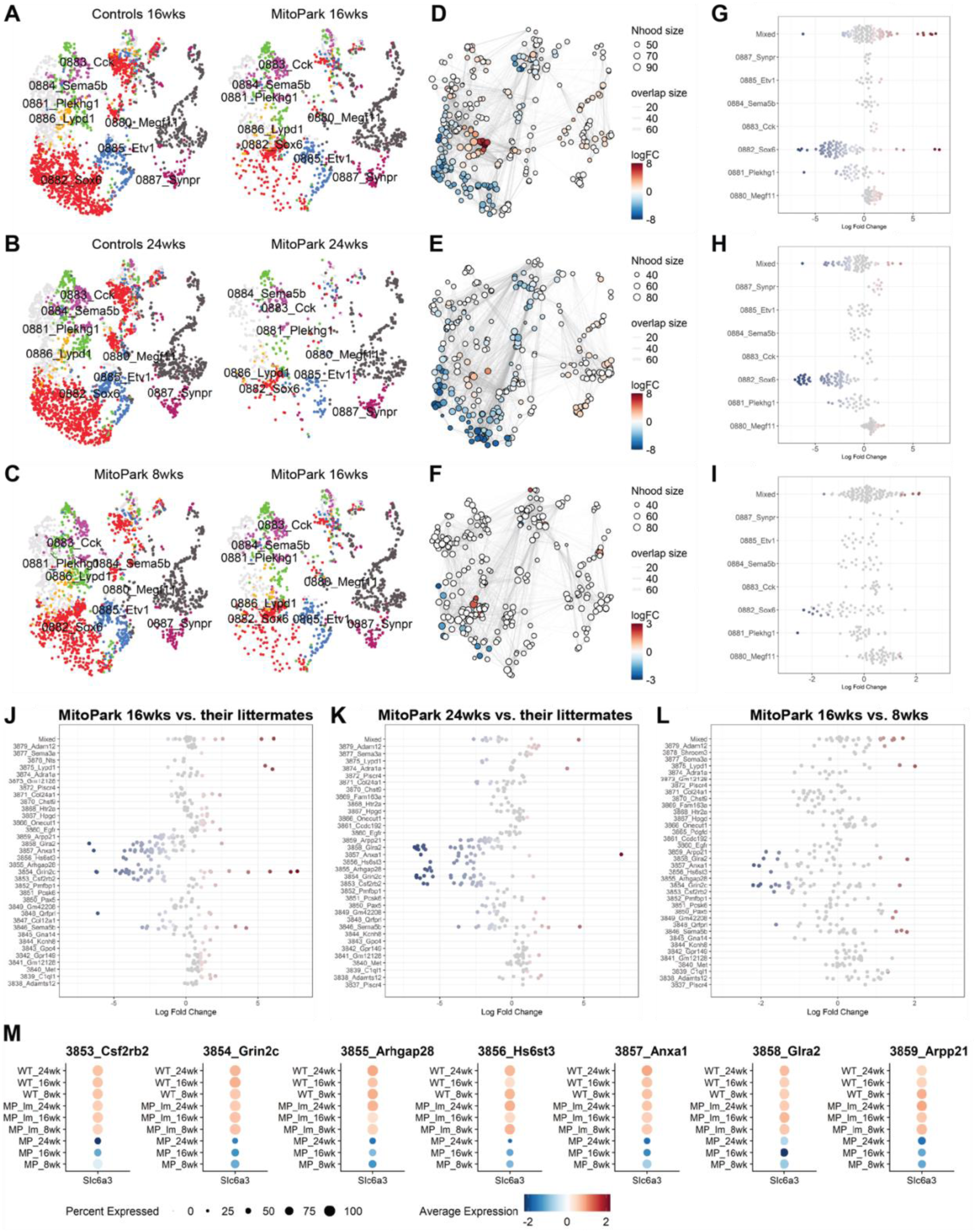
Selective vulnerability of *Sox6+* dopaminergic population revealed by differential abundance test with Milo, Related to Fig. 4. Differential abundance testing for the dopamine subpopulations with supertype annotations by the ABC atlas. **A-C.** UMAP plots showing the dopamine subpopulations of MitoPark mice and their controls. **D-F.** Neighborhood graphs with the results from Milo differential abundance analysis between MitoPark mice and their littermate controls (D:16 weeks, E: 24 weeks), and between MitoPark mice at different stage (F: 8 weeks and 16 weeks). **G-I**. Beeswarm plots showing the distribution of log-fold changes in each supertype group. Colors are represented similarly to **D-F**. **J-L**. Beeswarm plots showing the distribution of log-fold changes in each cluster. Colors are represented similarly to **Fig. 4D-F**. **M**. Comparison of DAT expression among *Sox6+* dopamine subtypes revealed no substantial differences.

**Extended Data Fig. 8.**
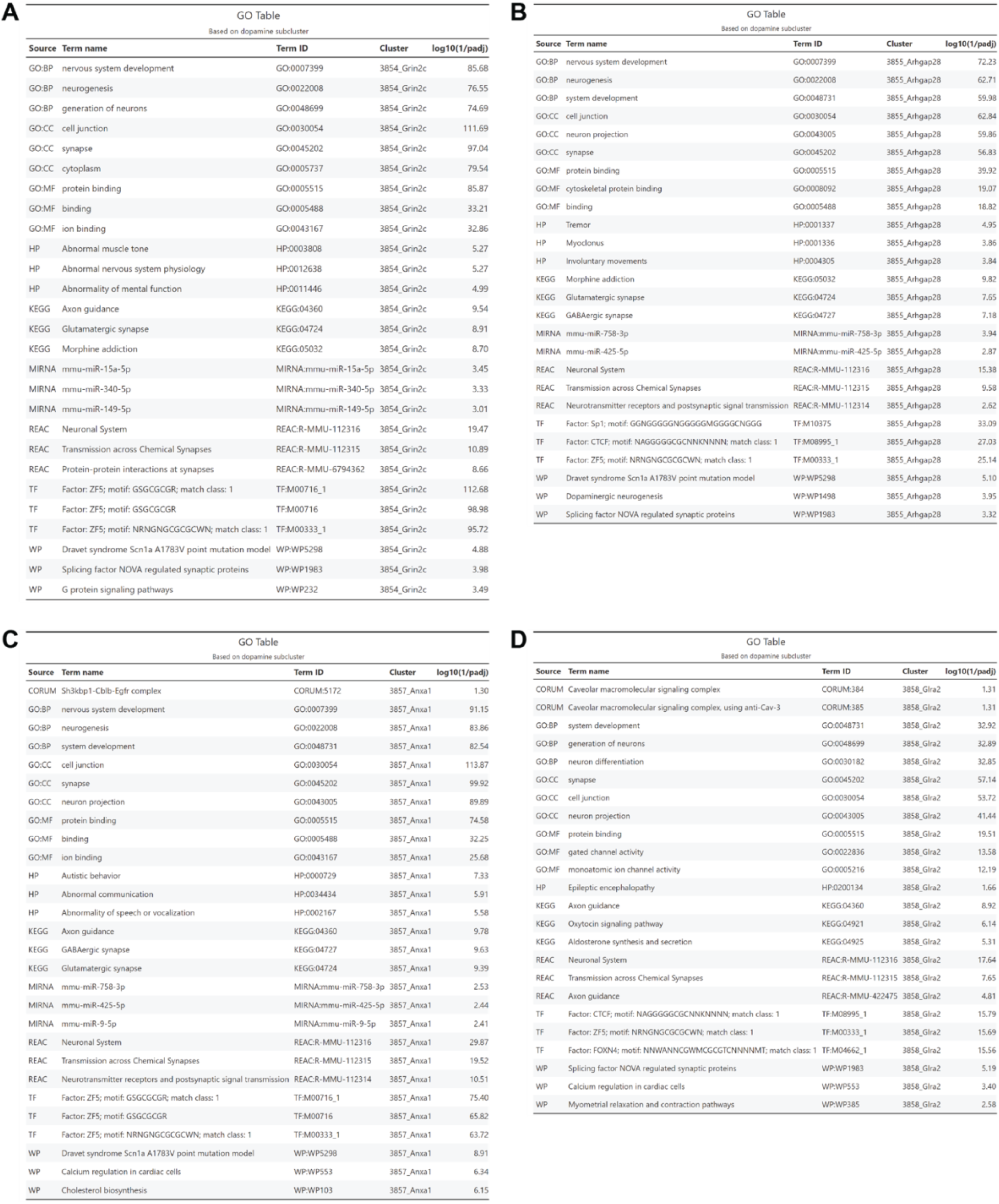
Gene Ontology (GO) enrichment analysis for vulnerable dopaminergic population, Related to Fig. 4. Enrichment analysis of gene ontology (GO) terms for key genes in a representative vulnerable dopamine cluster (A, 3854_Grin2c; B, 3855_Arhgap28; C, 3857_Anxa1; D, 3858_Glra2).

**Extended Data Fig. 9.**
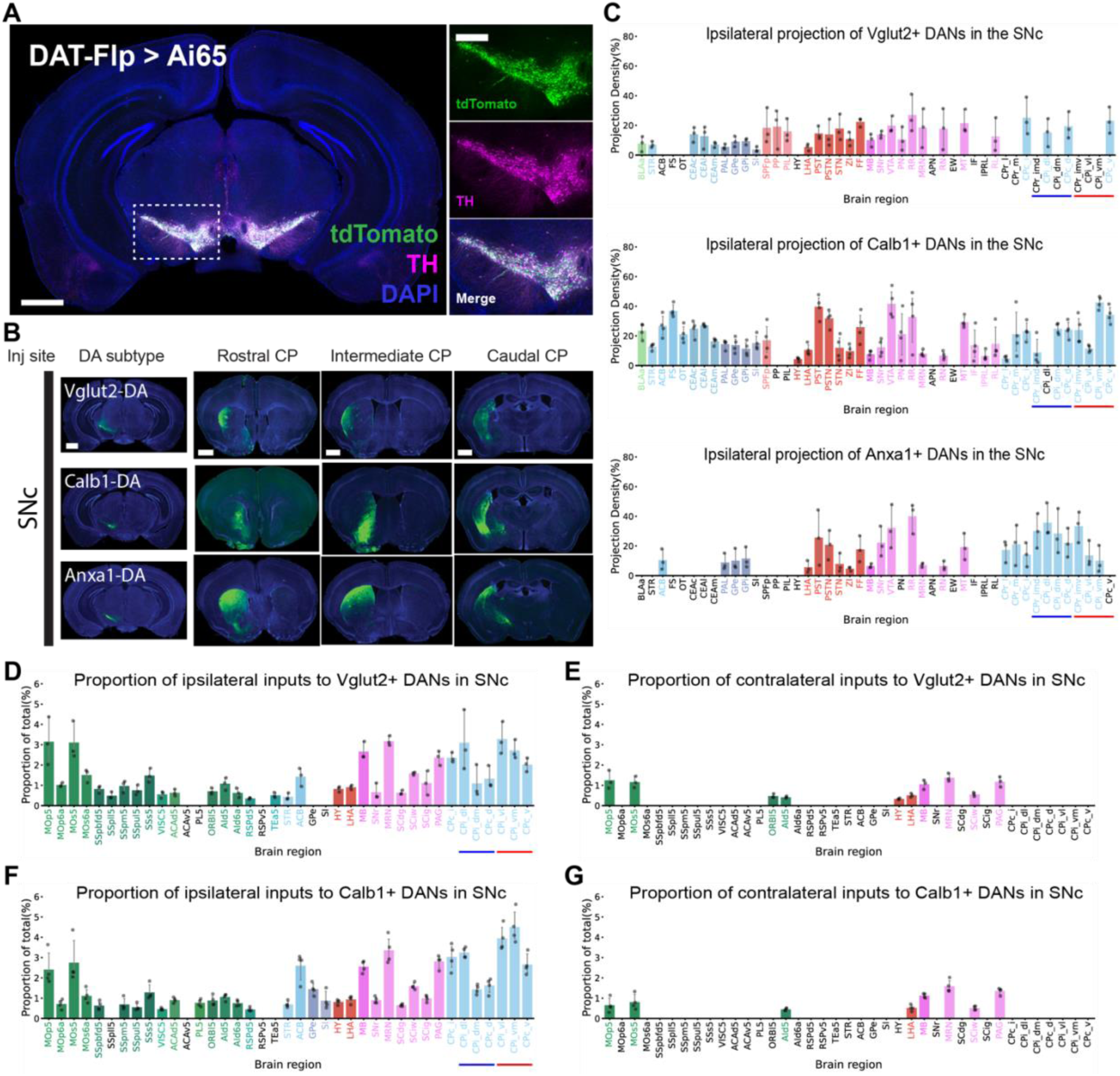
Whole-brain efferent and afferent connectivity of dopaminergic neuron subtypes, Related to Fig. 5. **A.** Characterization of the DAT-Flp line using a reporter line (Ai65) confirmed that tdTomato+ cells are *Th+* neurons. Scale bars: 1mm for the striatum (left) and 400μm for the substantia nigra (right). **B.** Representative images of axonal projections of *Vglut2+*, *Calb1+* and *Anxa1+* dopamine neuron subtypes in the SNc. They preferentially project to dorsolateral striatum, ventral medial striatum and dorsal striatum. Scale bar 1mm. **C.** Brain regions to which *Vglut2+*, *Calb1+* and *Anxa1+* dopamine neuron project, measured as the fraction of neurites found within those brain structures defined Allen brain atlas (n=4 *Calb1+* and n=3 for *Vglut2+* and *Anxa1+*). Blue horizontal line indicates dorsal striatum and red line indicates ventral striatum. **D-G.** Statistical analysis of the whole-brain distribution of ipsilateral (left) or contralateral (right) monosynaptic inputs to *Vglut2+* neurons (D, E) or *Calb1+* dopamine neuron subtype (F, G) in the SNc. Brain areas are color-coded by the Allen Brain Atlas.

**Extended Data Fig. 10.**
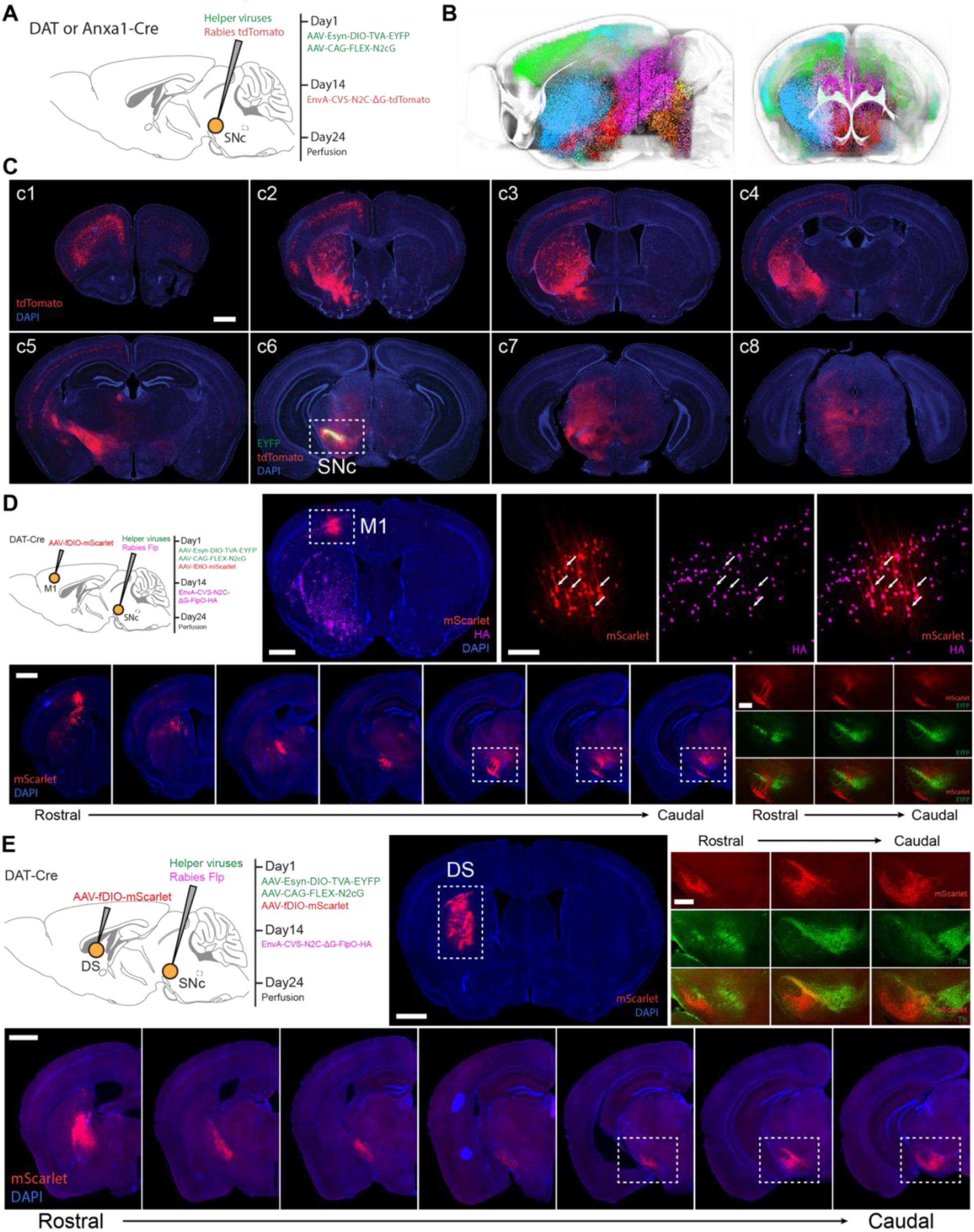
Direct afferent inputs from the motor cortex or striatum to the SNc DANs, Related to Fig. 5. **A.** Schematic of the rabies injection procedure and experimental timeline for helper viruses and rabies viruses in the DAT-Cre or Anxa1-Cre. **B.** Whole-brain 3D reconstruction using BrainJ, color coded by brain regions^19^ (left: sagittal view, right: coronal view). **C.** Representative images of monosynaptic inputs to DANs from the whole brain (c1-c8, from the rostral to caudal). The small white rectangle indicates the injection site, the SNc. Scale bar 1mm. **D-E.** Monosynaptic projections from primary inputs include cells associated with motor function to the SNc DANs (D, motor cortex; E, striatum). Scale bars: 1mm for the striatum and 400μm for the substantia nigra.

**Extended Data Fig. 11.**
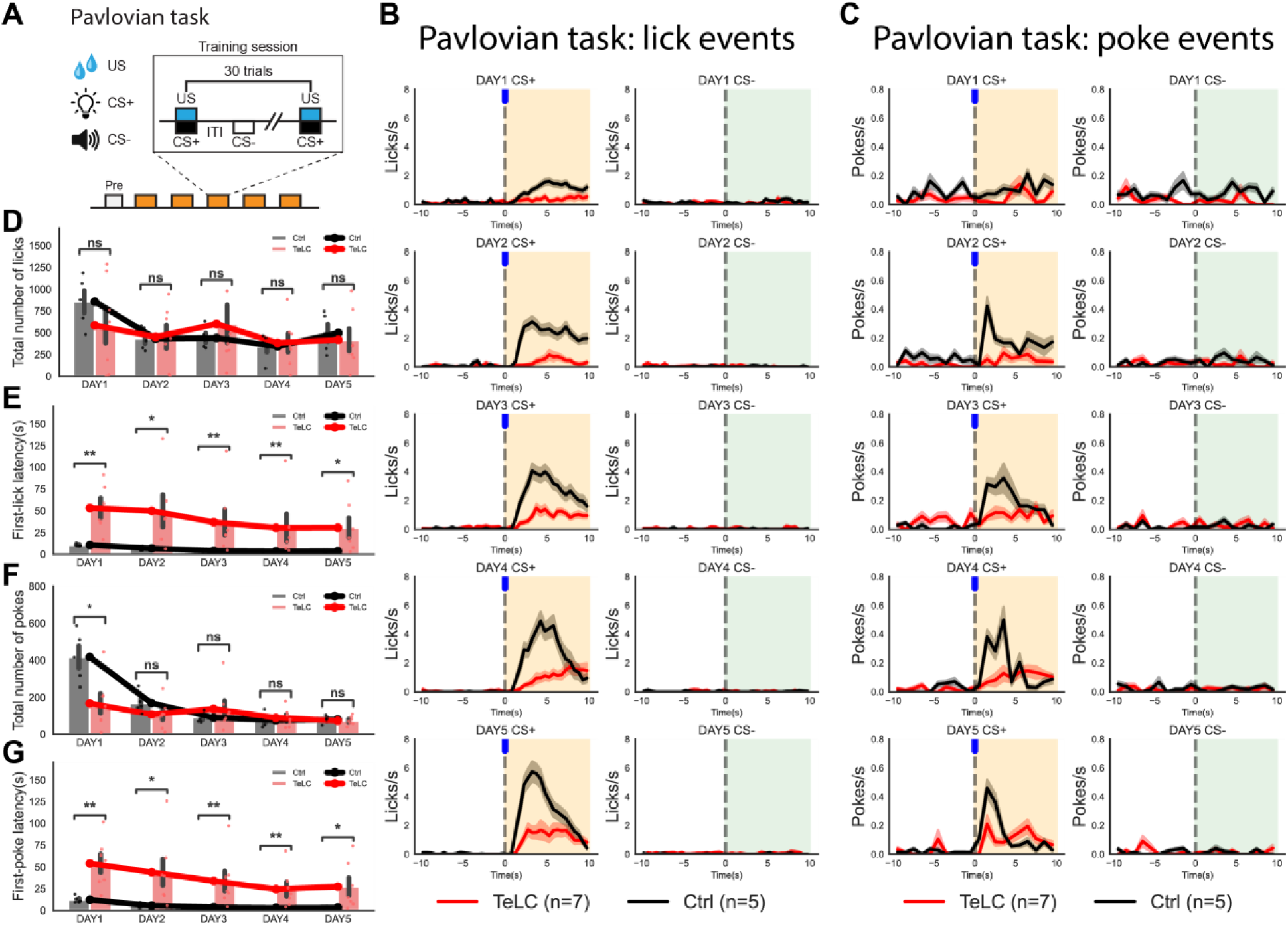
Related to Fig. 6. **A.** Schematic representation of the Pavlovian task. A light and a tone are used as CS+ and CS-respectively. **B-C.** PSTH (the peristimulus time histogram) of the lick or poke rate frequency of Anxa1-TeLC animals (red) and their controls (black) in response to CS+ (left) or CS- (right). The control animals consistently demonstrated a robust response to the light stimulus over time; in contrast, the Anxa1-TeLC animals exhibited a diminished response. D-G. Although the total number of licks did not differ significantly (D), there was a notable difference in the latency to the first lick between the Anxa1-TeLC and control animals (E). I-J. On Day 1, a difference in the number of pokes was observed, but this disparity did not persist in subsequent days (F). Additionally, there was a significant difference in the latency to the first poke between the mutant and control animals (G).

